# Two distinct oscillatory auxin signals define the plasticity of lateral rooting in *Arabidopsis thaliana*

**DOI:** 10.1101/2024.08.16.608248

**Authors:** Chengzhi Ren, Jule Bodendorf, Miguel A. Moreno-Risueno, Jürgen Kleine-Vehn

## Abstract

The phytohormone auxin defines lateral root pre-branch sites (PBS) in the growing primary root tip. How PBS contribute to the plasticity of the root system architecture remains incompletely understood^1,2,3,4^. Here, we reveal in the model plant *Arabidopsis thaliana* that two distinct oscillatory systems for the phytohormone auxin coordinate the spatial and temporal identity of PBS, jointly defining the lateral root density. We followed auxin signalling dynamics for days and thereby detected a systemic auxin signal oscillating along the mature primary root, where PBS potentially form functional root branches. While a previously proposed local oscillatory auxin zone spatially primes the PBS in the growing root tip, we reveal that the systemic oscillatory auxin signal temporally controls the auxin-dependent identity of these PBS. Light perception in the shoot defines the strength of the systemic auxin signal and thereby controls the auxin-reliant ability of PBS to develop into lateral roots. Moreover, PHYB and CRY1 mediate the light-dependent integration of other environmental signals, such as ambient temperature, into the control systemic auxin signalling and lateral root density. Our work reveals how two spatially distinct oscillatory auxin signals define the plasticity of plant root development in response to fluctuating conditions.

## Main

*De novo* organogenesis is a hallmark of the remarkable plasticity and adaptability in plants. In the context of root system architecture, lateral root (LR) formation is a critical process that enhances nutrient and water uptake, providing stability and resilience. Although several models have been proposed, the mechanism of lateral root initiation, especially during the early stage, remains an active area of research. A hypothesis suggests that the sites of presumptive LR development are predetermined along the primary root axis through a mechanism which involves the so-called "root clock"^1,2,3,4^. According to this, a localized oscillatory signal of the phytohormone auxin in the elongation zone of the main root (oscillatory zone) primes cells to become the LR pre-branch sites (PBS)^2^. Although the root clock model can describe the regular formation of PBS, there remain open questions regarding an apparent lack of plasticity in this system and the extent to which PBS are predetermined to ultimately develop into LR^5^.

We used prolonged imaging of the synthetic, auxin-responsive *DR5* promoter fused to a luciferase reporter (*pDR5::Luc*)^2^ to visualize the nuclear auxin output signalling and therewith PBS dynamics for days at constant light (120 μmol·m^-2^·s^-1^) and temperature (21°C) conditions. Therewith, we revealed that a substantial fraction of the PBS lost the auxin signal in time and did not give rise to LR development (Supplementary Fig. 1). We, hence, addressed the currently elusive mechanism that decisively defines the transient or persistent nature of the auxin-reliant PBS and its contribution to control LR density.

### Light quantitatively controls a systemic oscillatory auxin signal

To analyze the long-term dynamics of *DR5* luciferase activity, we utilize kymograph representations, which are graphical depictions of spatial position (top to bottom) over time (left to right), essentially capturing the dynamic changes along a root in a single image. This approach allowed us to detect a repeated fluctuation of auxin signalling over time. These dynamics were present along the primary root, including the PBS, suggesting a systemic synchronization (Fig. 1a,b and Supplementary Fig. 2a; Supplementary Fig. 3). In our constant growth condition, the frequency of this systemic auxin signalling fluctuation followed a normal distribution with a period of around 24 hours (Fig. 1c, Supplementary Fig. 2b). We accordingly define these dynamics hereafter as a systemic oscillatory auxin signal. We subsequently assessed if environmental factors, such as light intensity, affect this systemic oscillatory auxin signal. When compared to control 120 μmol·m^-2^·s^-1^ light conditions (hereafter referred to as comparatively high light, HL), a reduction of the light intensity to 50 μmol·m^-2^²·s^-1^ ¹ (hereafter referred to as comparatively medium light, ML) did not affect the frequency of the systemic auxin oscillations (Fig. 1d,e and Supplementary Fig. 2c,d) but did quantitatively decrease the overall intensity of the systemic auxin signal (Fig. 1f). Notably, the systemic auxin signalling was undetectable at a light intensity of 20 μmol·m^-2^²·s^-1^¹ (hereafter referred to as comparatively low light, LL) (Fig. 1g and Supplementary Fig. 4).

**Figure 1.**
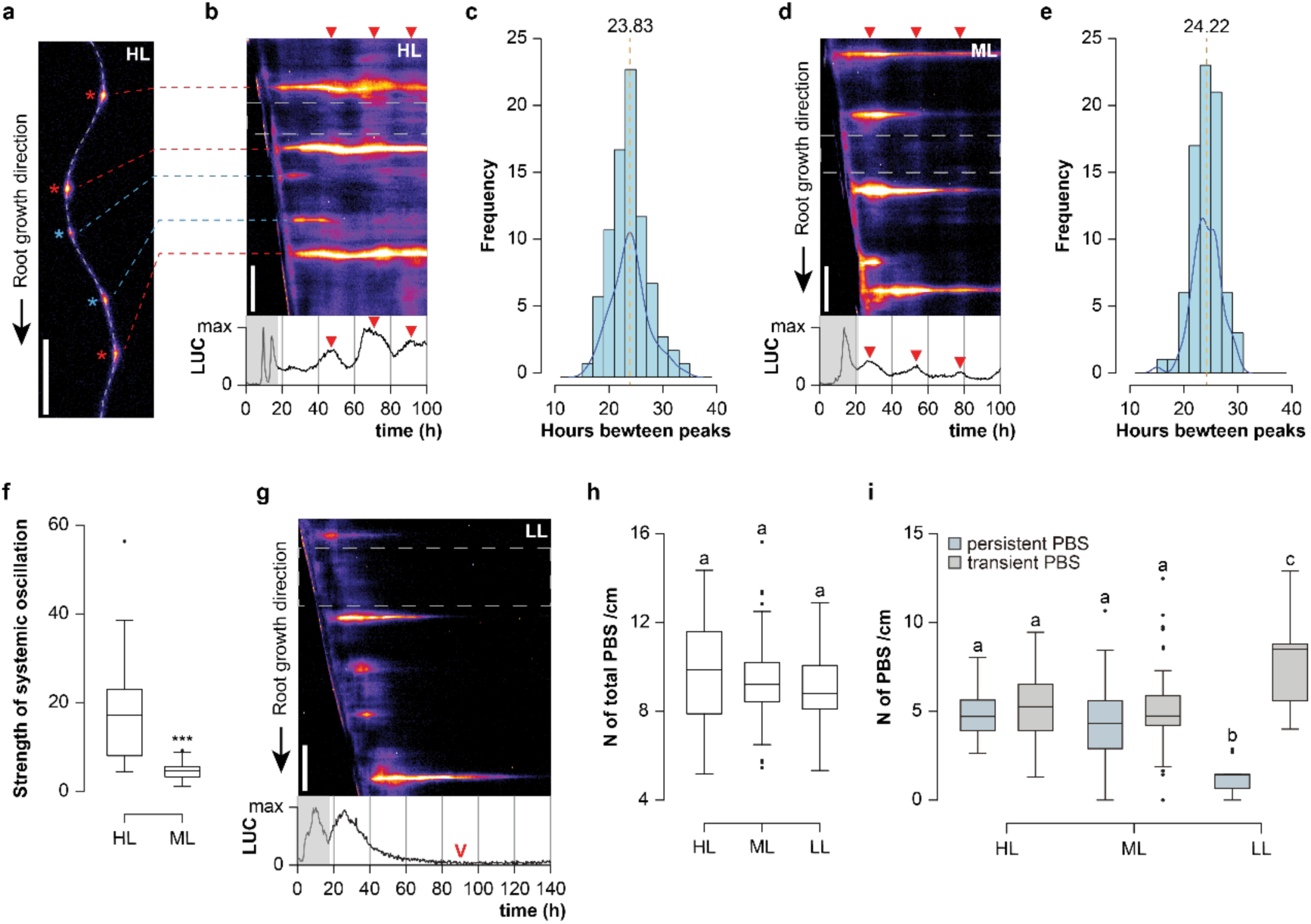
Light quantitatively defines a systemic auxin oscillation in main roots and pre-branch sites. **a-g,** *pDR5::Luc* expressing root (**a**) was segmented (dashed line) for the respective kymograph (**b**). Asterisks and dashed lines mark the positions of persistent (red) and transient (blue) auxin signals in PBS. Kymographs display auxin signalling dynamics in high light (HL) (**b**), medium light (ML) (**d**) and low light (LL) (**g**) (Scale bar = 1 mm). Quantification of real-time luminescence is below the kymographs where the red triangles mark oscillatory peaks of systemic auxin signal (**b**, **d**). Red arrowhead denotes the downregulation of the systemic auxin signal (**g**). The dashed grey rectangle in the kymographs mark the region for the real-time plots (**b**, **d**, **g**). The shaded area in real-time plots marked the signal originating from the root tip and root clock zone (**b**, **d**, **g**). Quantified distribution of the time intervals between oscillatory peaks of systemic auxin under (**c**) HL and (**e**) ML conditions (mean value marked by a yellow dashed line; *n* = 23 to 29; Anderson-Darling normality test, P-value = 0.4501 (**c**) and 0.177 (**e**)). (**f**) Quantified luminescence intensity of oscillatory peaks under HL and LL conditions (*n* = 20). **h**, Quantification of PBS density under HL, ML and LL conditions (*n* = 16 to 66). **i**, Quantification of persistent and transient auxin in PBS under HL, ML and LL conditions (*n* = 16 to 66). Paired and two-tailed student’s t-test performed for (**f**) (*P <* 0.001***). Letters indicate values with statistically significant differences from one-way ANOVA performed for (**h**) (*P* > 0.05) and (**i**) (*P <* 0.0001).

Based on these results, we propose that light quantitatively controls a systemic oscillatory auxin signal. Next, we addressed whether the systemic auxin signalling dynamics contribute to the transient nature of auxin-reliant PBS. Notably, despite the varying strength of systemic auxin signalling at different light intensities, we observed similar quantities of established PBS under HL, ML, and LL conditions (Fig. 1h), indicating that the initial priming of PBS is largely independent of light intensity. However, we observed a correlation between the impediment of systemic auxin signalling and a steep increase in transient auxin signal in PBS at LL conditions (Fig. 1i). This result suggests that light intensity contributes to maintaining the auxin-reliant identity of PBS, correlating with a quantitative impact on systemic auxin oscillation.

### Light-reliant systemic auxin signal oscillations define PBS identity

To specifically address the light dependency of already formed PBS, we monitored PBS formation in 5-day-old *pDR5::Luc* seedlings under HL conditions for 48 hours and afterwards exposed them either to LL conditions or maintained them under HL as a control. After transferring seedlings from HL-to-LL conditions, the systemic auxin signalling completely ceased, observing the last oscillatory peak after 8 hours (Fig. 2a and Supplementary Fig. 5a). Additionally, although the total quantity of PBS was similar under both light conditions (Supplementary Fig. 6), the HL-to-LL transfer revealed that light intensity is critical to maintaining auxin signalling in PBS when compared to seedlings kept in HL (Fig. 2c,d). These findings confirm that light intensity is crucial to maintaining the systemic oscillatory auxin signal and PBS identity.

**Figure 2.**
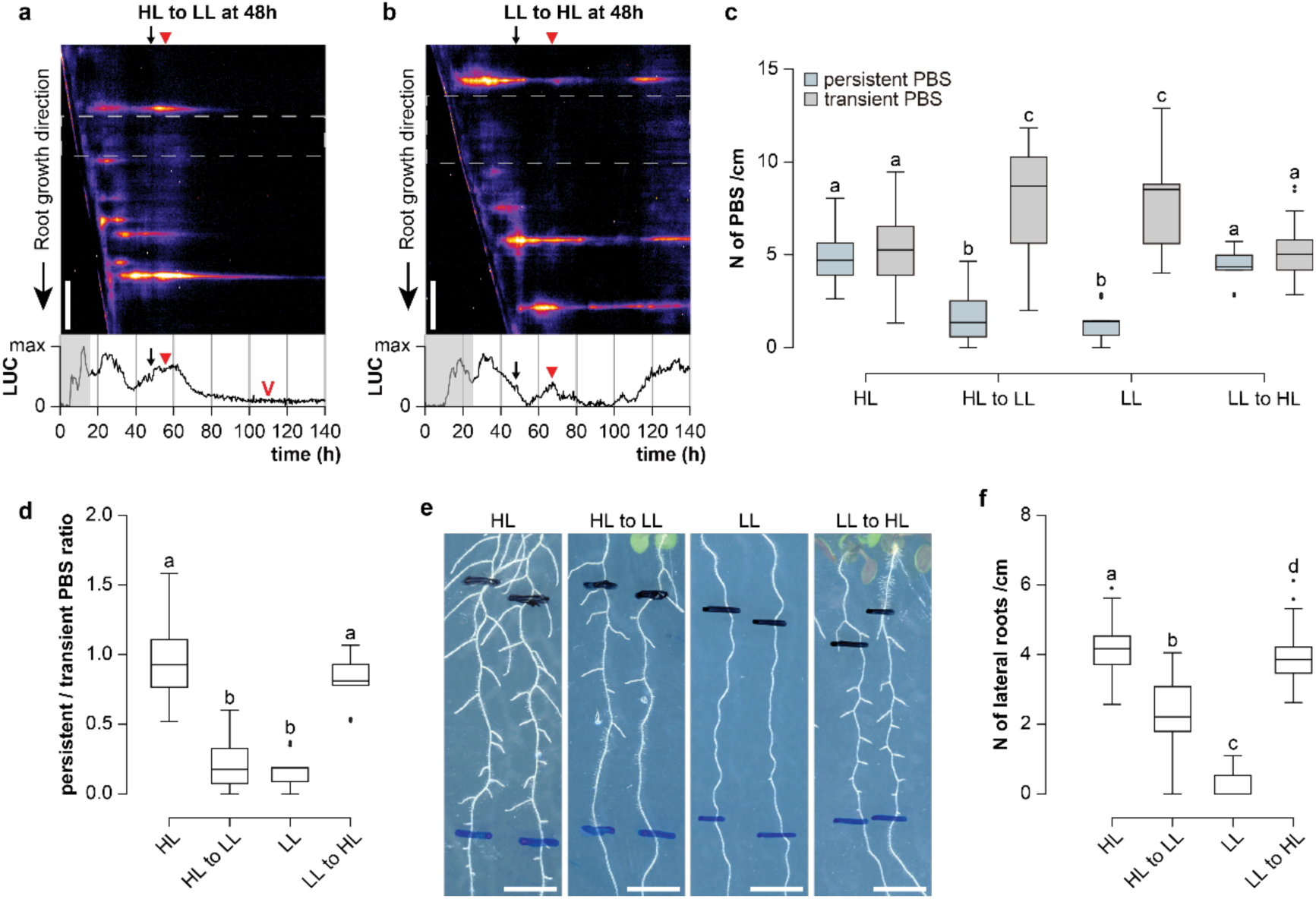
Light-reliant systemic auxin signal determines PBS dynamics and lateral rooting. **a-d**, Kymograph and real-time quantification of relative luminescence show systemic auxin and PBS progression dynamics during light transitions from high light (HL) to low light (LL) (**a**) and LL-to-HL (**b**) (Scale bar = 1 mm). Black arrows mark the time of transfer (**a**, **b**). The red triangle marks the peak of the systemic auxin signal (**a**, **b**). The red arrowhead indicates that auxin signalling has reduced to the background level(**a**). The dashed grey rectangle marks the region for the real-time plots; The shaded area in real-time plots marked the signal originating from the root tip and root clock zone (**a**, **b**). Quantification (**c**) and the corresponding ratio (**d**) of persistent and transient auxin signal in PBS under both transitional and maintained light conditions (*n* = 16 to 38). **e**, **f**, Representative images of root systems (**e**) and quantification of LR density (**f**) under both maintained light conditions and transferred HL-to-LL and LL-to-HL (*n* = 58 to 61; Scale bar = 5 mm). Letters indicate values with statistically significant differences from one-way ANOVA performed for (**c**), (**d**) and (**f**) (*P <* 0.0001 for all).

Contrariwise, we also monitored PBS formation in 5-day-old *pDR5::Luc* seedlings under LL conditions for 48 hours and subsequently transferred the seedlings either to HL conditions or kept them at LL as a control. In this condition, systemic auxin dynamics recovered around 18 hours after transferring from LL-to-HL (Fig. 2b and Supplementary Fig. 5b). Moreover, HL increased the recovery of auxin signalling in PBS when compared to those kept in LL (Fig. 2c,d). This finding again pinpoints the positive impact of light on systemic auxin signalling and its ability to reactivate auxin-reliant PBS.

Next, we tested if the light-dependent modulation of auxin in PBS indeed contributes to the reshaping of the root system architecture. To ensure the light effects are independent of LR priming, we transferred 7-day-old seedlings between different light conditions and analyzed three days later the LR appearance in a defined section of the main root (grown between days 4 and 7; see marks in Figure 2e). We observed a strong decrease in LR density when transferring roots from HL-to-LL compared to those maintained at HL (Fig. 2e,f). Conversely, LR density significantly increased when transferring from LL-to-HL as compared to those kept at LL (Fig. 2e,f). This set of data confirms that light conditions control auxin-dependent PBS and thereby LR density. Accordingly, we propose that light quantitatively controls systemic oscillatory auxin signals, thereby temporally defining the identity of previously primed PBS and ultimately LR spacing.

### Systemic auxin signal controls PBS identity at high ambient temperature

We hypothesize that while the auxin oscillation zone enables the regular priming of PBS during main root growth, a systemic oscillatory auxin signal could integrate various environmental information into lateral root spacing. To further test this assumption, we introduced variation in ambient temperature. Temperature and light signalling both play a crucial role in shaping root system architecture and are in part, molecularly interconnected^6^. Nonetheless, the existing literature reveals inconsistent findings, suggesting that elevated ambient temperatures can exert both positive and negative effects on lateral rooting within and across different species^7,8,9,10,11,12,13^. To shed light on this seemingly complex role, we initially tested whether temperature also affects auxin signalling dynamics in the root, we transferred 5-day-old *pDR5::Luc* seedlings from a control temperature of 21°C to a high ambient temperature (HT) of 29°C at constant HL condition. By monitoring the root luminescence after the transfer, we observed an overall increase of auxin signalling in the oscillation zone corresponding with an increased signal in the kymograph diagonal (Fig. 3a,b), which also correlated with an increased number of PBS (Fig. 3a,c). However, the vast majority of auxin-reliant PBS appeared transiently, aligning with a severe systemic disruption of auxin signalling over time (Fig. 3d,e, Supplementary Fig. 7).

**Figure 3.**
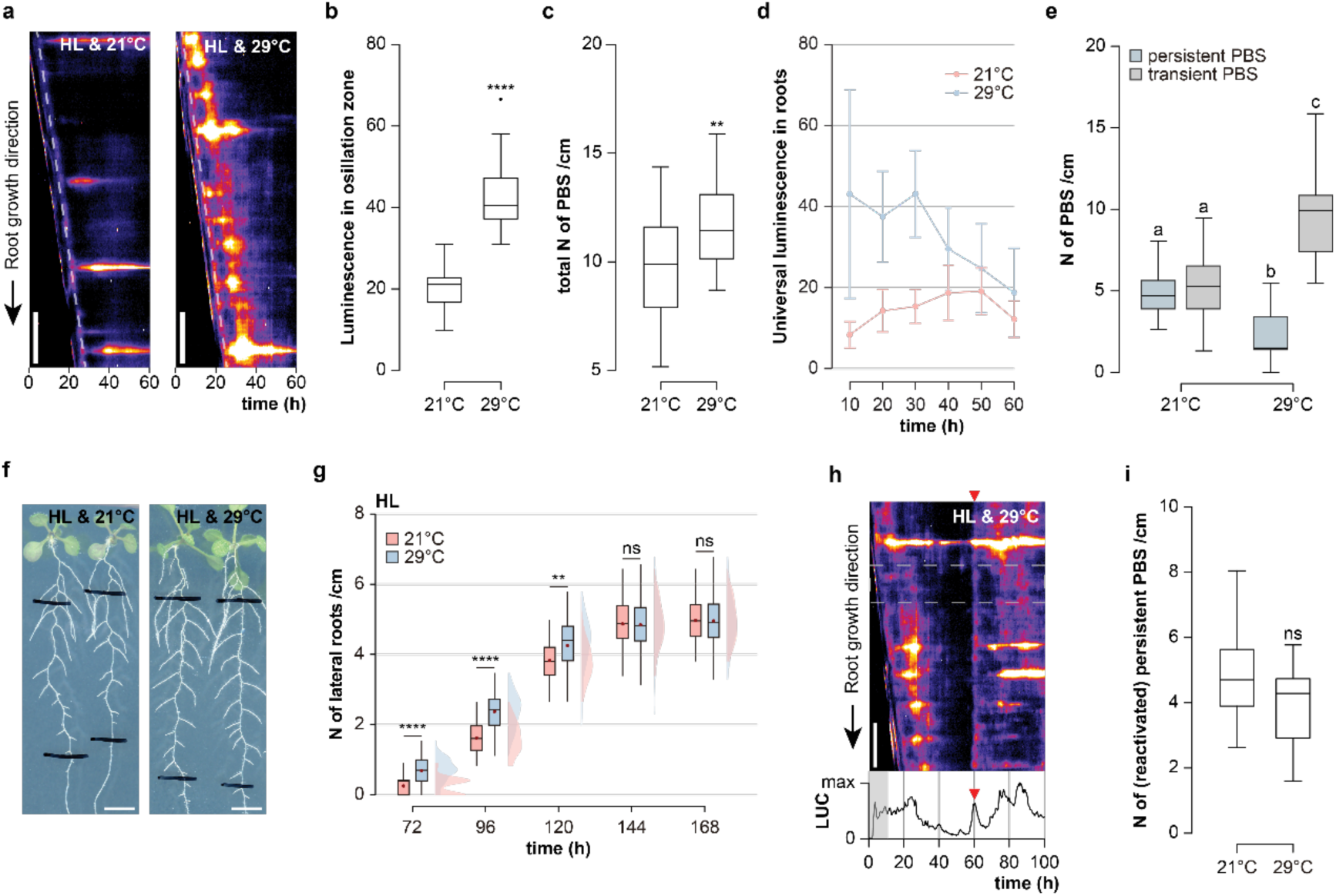
High-temperature affects systemic auxin signal and PBS dynamics. **a**, Kymograph of *pDR5::Luc* expressing roots, showing the priming and progression of PBS at 21°C and 29°C under high light (HL) conditions. The dashed line marks the local oscillatory (root clock) zone (Scale bar = 1 mm). **b**, Quantification of average luminescence in oscillatory zone over time in 21°C and 29°C under HL conditions (*n* = 24 to 30). **c**, Quantification of PBS density at 21°C and 29°C under HL conditions (*n* = 16 to 22). **d**, The quantified dynamic of *pDR5::Luc* signal in the main roots at 21°C and 29°C under HL conditions (Error bars represent standard deviation; *n* = 16 to 22). **e**, Quantification of persistent and transient PBS at 21°C and 29°C under HL conditions (*n* = 16 to 22). **f**, **g**, Representative images of the root system (**f**) and time series quantification of LR density (**g**) at 21°C and 29°C under HL conditions (*n* = 54 to 55; Scale bar = 5 mm). **h**, Long-term kymographs and real-time quantification of relative luminescence showing the recovery of systemic auxin and PBS under 29°C and HL condition. The red triangle marked the recovery peak of systemic auxin signal; The dashed grey rectangle marks the region for the real-time plots; The shaded area in real-time plots marked the signal originating from the root tip and root clock zone (Scale bar = 1 mm). **i**, Quantified density of persistent PBS at 21°C and reactivated persistent PBS at 29°C under HL condition during prolonged monitoring (*n* = 16 to 22). Paired and two tailed student’s t-test performed for (**b**), (**c**), (**g**) and (**i**) (*P <* 0.01** and < 0.0001****). Letters indicate values with statistically significant differences from one-way ANOVA performed for (**e**) (*P <* 0.0001).

Our results, hence, suggest that HT initially imposes a positive effect on initiating PBS but subsequently limits the systemic auxin signal along the primary root. Contrary to this observation, the lateral root primordia (LRP) (Supplementary Fig. 8) and overall LR density (Fig. 3f,g) were not inhibited by HT but showed even slight enhancement when compared to the control. To address this apparent contradiction, we initially assessed if the systemic reduction of *pDR5::Luc* at higher temperatures could relate to reduced enzymatic activity of luciferase at higher temperature. The reported midpoint of the unfolding transition temperature of the luciferase enzyme *in vitro* is around 34°C, which is further stabilized by osmolytes^14^. In agreement, luciferase activity in *p35S::Luc* expression control lines did not decrease when transferred to 29 °C (Supplementary Fig. 9). Notably, *p35S::Luc* showed at both temperatures an enhancement over time, and the signal was consistently higher at 29 °C, which may relate to variable 35S promoter activity in response to environmental changes or developmental cues^15^. We conclude that the systemic decrease in *pDR5::Luc* dynamics at high temperatures is not due to changes in enzymatic luciferase activity.

To further address the paradox of auxin dynamics and root system architecture, we extended our imaging of the auxin output signalling to 8 days. Thereby, we observed a recovery of the systemic auxin signalling around 70 hours after transferring seedlings to HT conditions (Fig. 3h and Supplementary Fig. 10). The systemic peak in auxin correlates with the reactivation of PBS, ultimately resulting in an increase in persistent PBS at HT (Fig. 3h,i). Notably, the number of recovered, persistent PBS under this condition is remarkably similar to the number in the control temperature condition (Fig. 3i). The reactivation of PBS strictly occurs simultaneously or after recovery of systemic auxin peak (Fig. 3h, Supplementary Fig. 10), suggesting again that a systemic auxin signal enables the integration of environmental cues into the temporal control of already primed PBS.

### Light perception modulates high temperature-dependent control of auxin-reliant PBS identity

To further address that the systemic auxin signal indeed integrates environment information into lateral rooting, we quantitatively lowered the HT-induced recovery of the systemic auxin signal, using medium light conditions. Accordingly, we grew *pDR5::Luc* seedlings at 21°C and HL and transferred 5-day-old seedlings to HT and ML conditions. Similar to seedlings at HL (Fig. 3), HT also increased the auxin signalling output at the local oscillation zone (Fig. 4a,b), correlating with initially enhanced PBS priming at ML conditions when compared to the control (Fig. 4c). Comparable to the HL condition, the formed PBS showed transient auxin signals, being reflected by a strong, systemic disruption of auxin signalling in the mature root.

**Figure 4.**
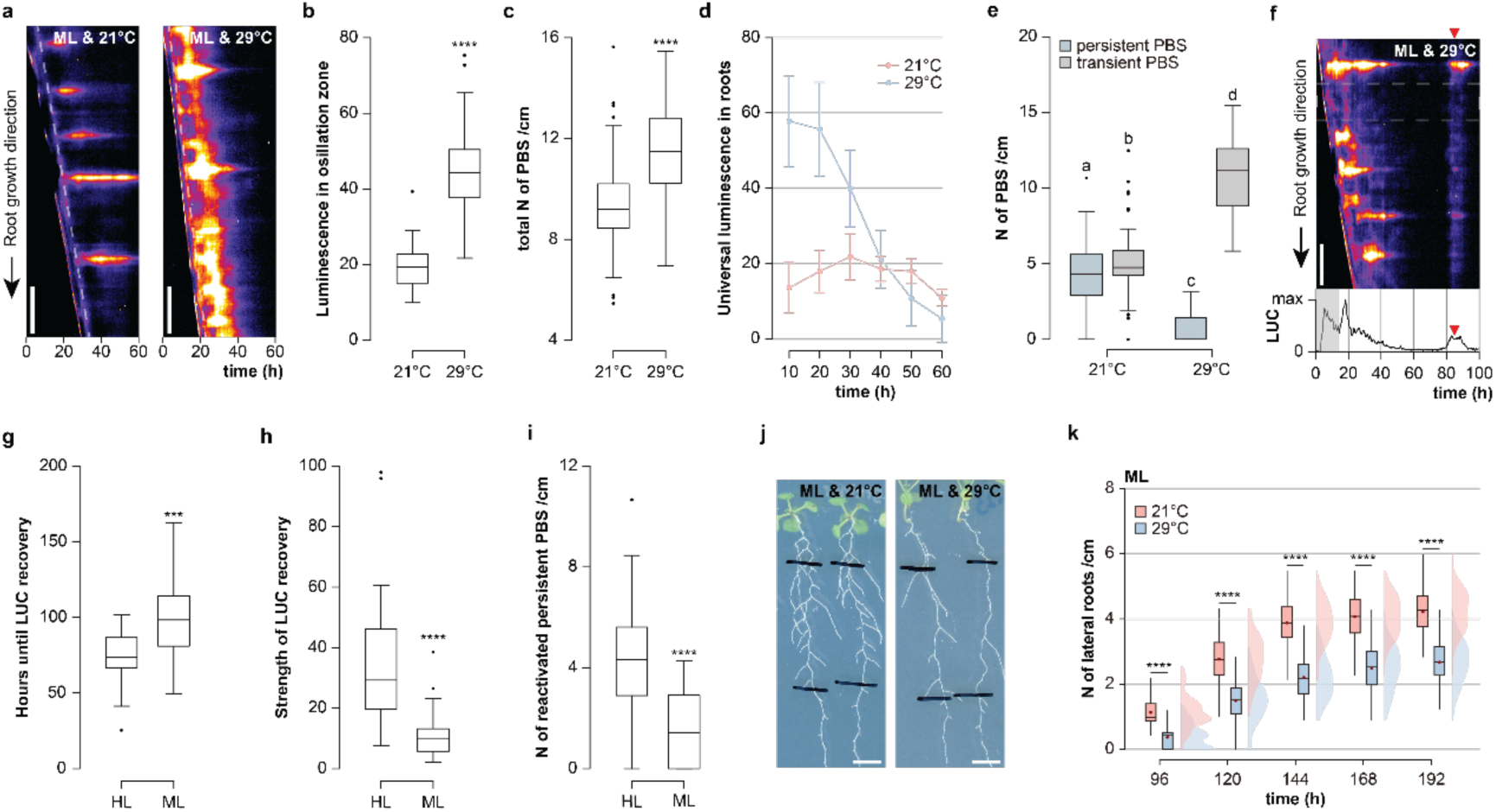
Light modulates high-temperature-dependent impact on systemic auxin signal and thereby lateral rooting. **a**, Kymograph of *pDR5::Luc* expressing roots, showing the priming and progression of PBS at 21°C and 29°C under medium light (ML) conditions. The dashed line marks the local oscillatory (root clock) zone (Scale bar = 1 mm). **b**, Quantifying average luminescence in oscillatory zone over time at 21°C and 29°C under ML conditions (*n* = 24 to 40). **c**, Quantification of PBS density at 21°C and 29°C under ML conditions (*n* = 66). **d**, Quantified dynamic of universal *pDR5::Luc* in main roots at 21°C and 29°C under ML conditions (Error bars represent standard deviation; *n* = 16 to 32). **e**, Quantification of persistent and transient PBS at 21°C and 29°C under ML conditions (*n* = 66). **f**, Long-term kymographs and real-time quantification of relative luminescence showing the recovery of systemic auxin and PBS at 29°C under ML condition. The red triangle marks the recovery peak of the systemic auxin signal; The dashed grey rectangle marks the region for the real-time plots; The shaded area in real-time plots marks the signal originating from the root tip and local oscillation (root clock) zone (Scale bar = 1 mm). **g**, **h**, Quantification of time until recovery (**g**) and strength (**h**) of systemic auxin signalling peak in primary roots after transferred at 29°C under ML and high light (HL) conditions (*n* = 24 to 48). **i**, Quantified density of reactivated persistent PBS at 29°C under ML and HL conditions (*n* = 24 to 48). **j-k**, Representative images of the root system (**j**) and time series quantification of LR density (**k**) at 21°C and 29°C under ML conditions (*n* = 54 to 55; Scale bar = 5 mm). Paired and two-tailed student’s t-test performed for (**b**), (**c**), (**g**), (**h**), (**i**), (**k**) (*P <* 0.001*** and < 0.0001****). Letters indicate values with statistically significant differences from one-way ANOVA performed for (**e**) (*P <* 0.0001 for all).

The HT also induced a temporal decline (Fig. 4d,e and Supplementary Fig. 11) and systemic recovery of auxin signalling in ML conditions (Fig. 4f,g and Supplementary Fig. 12). Importantly, the recovery of the systemic signal was, as anticipated, quantitatively reduced (Fig. 4h) when compared to HL conditions. Notably, the systemic auxin recovery was also temporarily delayed, occurring around 100 hours after the transfer (Fig. 4f,g and Supplementary Fig. 12). In agreement with our assumptions, ML-dependent reduction in systemic auxin signal also reduced the recovery of auxin-reliant PBS (Fig. 4i) when compared to HL conditions. This set of data strongly supports that light-dependent, systemic auxin signalling defines the transient or persistent nature of PBS. Moreover, our data suggests that light quantity modulates the negative impact of HT on PBS progression.

We next tested whether the proposed light-dependent modulation of ambient temperature response indeed defines root system architecture. In contrast to the limited effect of HT on LR density under HL conditions (Fig. 3), HT induced a strong reduction in the density under ML conditions (Fig. 4j,k), confirming that light quantity indeed contextualizes the integration of ambient temperature into root system architecture.

Next, we tested if this contextualizing mechanism defines not only PBS identity but also blocks the LRP development. Whereas the LR density was decreased in these conditions, we observed a quantitively similar amount of LRP when compared to the control condition (Supplementary Fig. 13). This finding suggests that light-modulated HT response does not primarily block LRP progression but impacts on PBS identity. In agreement with its impediment in systemic auxin signalling, we observed that the HT-dependent inhibition of lateral rooting was further enhanced under LL conditions (Supplementary Fig. 14). This set of data suggests that systemic oscillation of auxin integrates environmental signals, such as light and temperature, into the temporal control of PBS, contributing significantly to the root system architecture. We hence propose that the effect of high ambient temperature on lateral rooting is conditional, which can explain the seemingly disparate literature on high temperature and its impact on lateral rooting^7,8,9,10,11,12,13^.

Besides its impact on lateral rooting, high temperature also enhances main root growth in an auxin-dependent manner^16,17,18^. Moreover, it has been proposed that the priming of lateral roots may depend on the interplay of auxin and main root growth dynamics^19,20^. Hence, HT could indirectly affect lateral rooting by impacting primary root growth. Therefore, we next address whether HT-induced main root growth is linked to lateral rooting under our conditions. HT induced main root growth in HL, but this effect was abolished in ML conditions, which is conversely to its impact on lateral rooting (Supplementary Fig. 15a,b). These findings indicate that HT affects lateral rooting and main root growth via distinct mechanisms.

### Oscillatory IAA14 coordinates systemic auxin signalling to modulate lateral root PBS

To further address the here proposed contextualizing role of auxin in light- and HT-dependent inhibition of lateral rooting, we transferred 4-day-old seedlings to 29°C or 21°C under ML conditions for 3 days and subsequently sprayed either with the endogenous auxin indole-3-acetic acid (IAA; 0.2 µM), auxin antagonist auxinole (20 µM), or Dimethyl sulfoxide (DMSO) as a solvent control. We analyzed lateral root density in these root sections after an additional 7 days of HT treatment. The application of auxin completely suppressed the HT-dependent inhibition of lateral rooting at ML condition (Fig. 5a-c). In contrast, auxinole-dependent inhibition of auxin signalling led to enhanced HT-induced inhibition of lateral root density compared to DMSO (Fig. 5a-c). These results confirm that HT-dependent inhibition of lateral rooting is linked to auxin responses.

**Figure 5.**
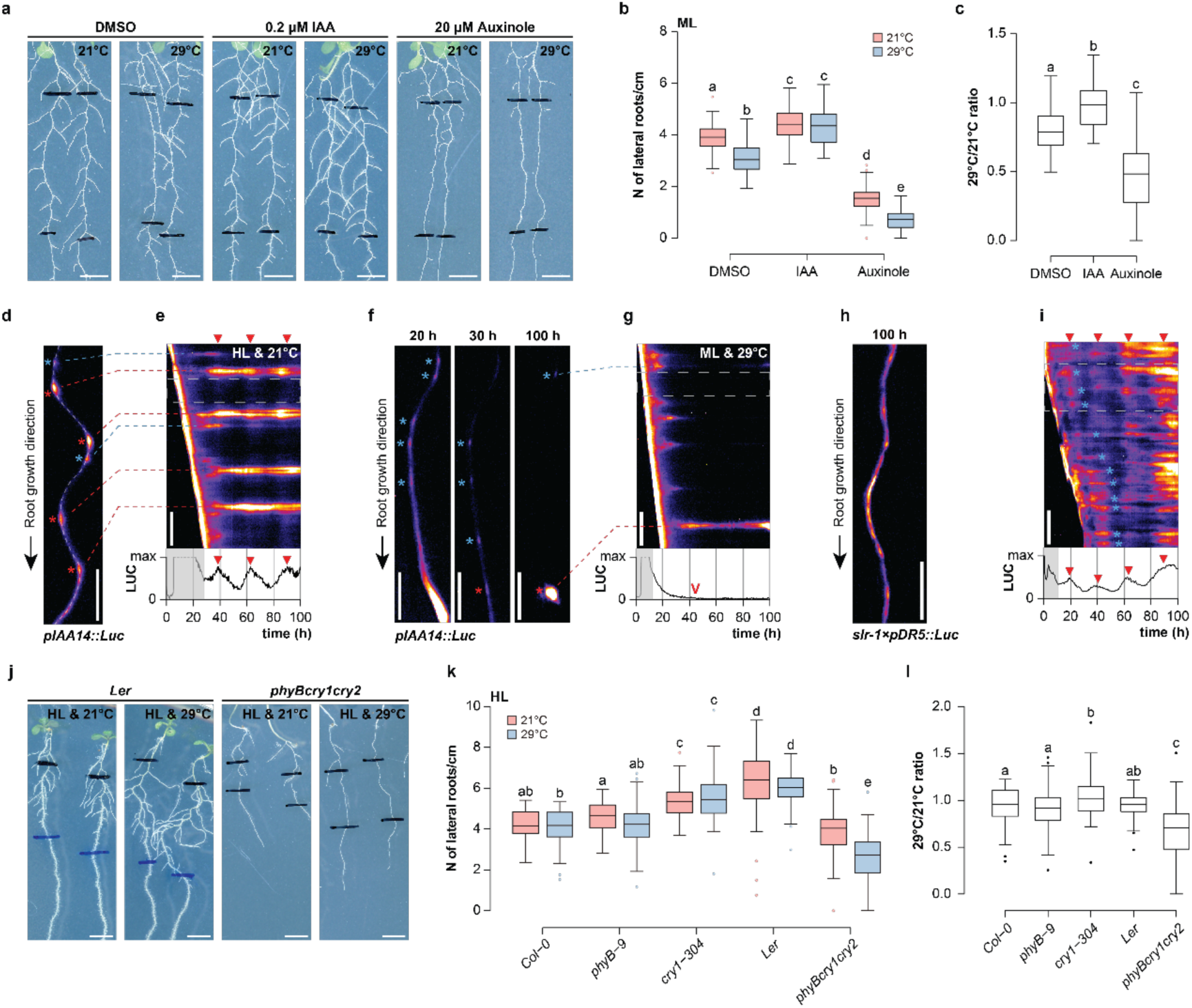
Dynamic of *IAA14* integrate systemic auxin signalling into the identity of PBS. **a**, **b**, Representative images of the root system (**a**) and quantification of LR density (**b**) at 21°C and 29°C under DMSO, 0.2 μM IAA and 20 μM Auxinole spraying (*n* = 55 to 63; Scale bar = 5 mm). **c**, Calculated ratio of final LR density at 21°C and 29°C spreading DMSO, 0.2 μM IAA and 20 μM Auxinole (*n* = 55 to 63). **d**-**g**, *pIAA14::Luc* expressing root at 21°C under high light (HL) (**d**) and 29°C under medium light (ML) (**f**). Asterisks and dashed lines (**d**, **f**) mark the positions of persistent (red) and transient (blue) PBS. Kymographs display *pIAA14::Luc* dynamics at 21°C under HL (**e**) and 29°C under ML (**g**) (Scale bar = 1 mm). Quantification of real-time luminescence is below the kymographs where the red triangles mark oscillatory peaks of *pIAA14::Luc* signal (**e**, **g**). Red arrowhead denotes the downregulation of the *pIAA14::Luc* signal (**g**). The dashed grey rectangle in the kymographs mark the region for the real-time plots (**e**, **g**). The shaded area in real-time plots marks the signal originating from the root tip and local oscillation (root clock) zone (**e**, **g**). **h**, **i**, *pDR5::Luc* expressing root (**h**) and respective kymograph (**i**) in *slr-1* background (Scale bar = 1 mm). Blue asterisks mark the positions of PBS-like pre-signals (**i**). Quantification of real-time luminescence is below the kymographs where the red triangles mark oscillatory peaks of auxin signal (**i**). The dashed grey rectangle in the kymographs mark the region for the real-time plots (**i**). The shaded area in real-time plots marks the signal originating from the root tip and local oscillation (root clock) zone (**i**). **j**, **k**, Representative images (**j**) and quantified LR density (**k**) of *Col-0*, *Ler* wild types and light receptor mutants at 21°C and 29°C under HL conditions (*n* = 45 to 72; Scale bar = 5 mm). **l**, Quantified ratio of LR density at 29°C to 21°C for *Col-0*, *Ler* wild types and light receptor mutants (*n* = 45 to 72). Letters indicate values with statistically significant differences from one-way ANOVA performed for (**b**), (**c**), (k) and (l) (*P <* 0.0001 for all).

We subsequently focused on SOLITARY ROOT/Indole-3-Acetic Acid inducible 14 (SLR/IAA14), a key repressor of nuclear auxin responses being involved in limiting lateral root initiation. The prominent gain-of-function *solitary root* (*slr-1*) mutant carries stabilized, auxin-insensitive *IAA14*, causing constitutive auxin repression and complete loss of lateral roots, which might be independent of PBS priming^21,22,23^.

To investigate the real-time transcription of *IAA14* in the root, we fused the promoter of *IAA14* to a luciferase reporter (*pIAA14::Luc*). By monitoring the luminescence in the root, we observed a continuous, non-oscillatory transcription of *IAA14* in the auxin oscillation zone of the root tip (Supplementary Fig. 16). This aligns with the hypothesis that *IAA14* operates independently of PBS priming, although an indirect effect cannot be disregarded. On the contrary, we observed systemic oscillatory dynamics of the *IAA14* reporter along the primary root and PBS, which is reminiscent of the synthetic auxin reporter *pDR5::Luc* dynamics under control conditions (Fig. 5d,e, Supplementary Fig. 17a). Additionally, The frequency of *IAA14* reporter fluctuation also followed a normal distribution with a period of around 24 hours (Supplementary Fig. 17a-c), similar to the oscillatory frequency of *pDR5::Luc* (Fig. 1c,e, Supplementary Fig. 2). Notably, *IAA14* reporter signal gradually became undetectable in primary root and PBS under LL condition (Supplementary Fig. 18), consistent with the disappearance of the *pDR5::Luc* signals (Fig. 1, Supplementary Fig. 4). These observations reaffirmed that light quantitatively modulates the systemic oscillatory auxin signalling. Similar to the synthetic *DR5* auxin signalling reporter, *IAA14* expression systemically decreased in both the primary root and PBS after transferring to HT under ML conditions (Fig. 5f,g). Unlike the strong systemic recovery of *pDR5::Luc* under HT (Fig. 3h, Fig. 4f, Supplementary Fig. 10 and Supplementary Fig. 12), *pIAA14::Luc* expression recovered specifically in the PBS (Fig. 5f,g). After transferring to HT and HL conditions, *IAA14* expression decreased exclusively in the primary root but remained persistent in individual PBS (Supplementary Fig. 19). This set of data suggests that temperature and light-dependent control of systemic auxin signalling defines IAA14-reliant lateral rooting.

Next, we genetically interfered with these auxin signalling dynamics by introducing the *slr-1* mutation into the *pDR5::Luc* background by genetic crossing. We detected a systemic oscillatory auxin signal along the primary root of the *slr-1* mutant (Fig. 5h,i). This suggests that SLR is not required to generate the systemic auxin signal, but functions a downstream effector regulating PBS capacity to develop LRs. In agreement with this assumption, the PBS failed to be temporarily maintained in *slr-1* between the systemic oscillation peaks of auxin (Fig. 5h,i). This set of data reinforces that the integration of systemic auxin signalling is critical for persistent PBS identity and its ability to develop into lateral roots.

### PHYB and CRY1 mediate shoot-perceived light modulating effect of high ambient temperature on lateral rooting

Our collective findings highlight the principal role of light in modulating temperature-dependent systemic auxin signalling and subsequent lateral rooting. We subsequently assessed where light is perceived to contextualize the temperature-sensitive lateral rooting. Therefore, we used light exposure of the root or shoot only to dissect where the light signal that modulates lateral rooting is perceived. We found that light exposure of the shoot is not only essential for lateral root development but also sufficient to modulate the temperature-dependent lateral rooting (Supplementary Fig. 20a-c). Hence, we assume that light perception in the shoot contributes to the auxin-reliant lateral rooting mechanism. Photosynthetic sucrose is an important determinant of lateral rooting^24^. However, the presence or absence of sucrose in the media did not affect the pattern of lateral rooting by HT under both HL and ML conditions (Supplementary Fig. 20d-f). This finding implies that photosynthetic sucrose is not the contextualizing signal in this context, pinpointing a light perception mechanism. Subsequently, we investigated the light perception mechanism of the light-modulated HT-dependent repression of lateral rooting. Mutations in the *phytochrome B* (*phyB*) red and far-red light photoreceptor caused hypersensitivity to HT-induced repression of lateral rooting at ML conditions when quantitatively compared to the wild-type (Supplementary Fig. 21a-c). In addition to functioning as a light sensor, PHYB also acts as a thermosensor^25^.

However, the increased hypersensitivity to HT-induced repression of lateral rooting observed in the *phyB-9* mutant suggests that PHYB does not function as the thermosensor in this process (Supplementary Fig. 21a-c). Moreover, we detected a similar hypersensitivity to HT at ML condition in mutants of the blue light receptor *cryptochrome 1* (*CRY1*) (Supplementary Fig. 21a-c). Notably, triple mutants of *phyBcry1cry2* not only enhanced hypersensitivity to HT-induced repression of lateral rooting at ML conditions compared to single mutants of *phyB-9* and *cry1-304* (Supplementary Fig. 21a-c), but also caused sensitivity to HT at HL conditions when compared to the wild-type, which was not observed in the single mutant of *phyB-9* and *cry1-304* (Fig. 5j-l). In contrast, the genetic interference with *phytochrome A* (*phyA*) or *phototropins* (*PHOTs*) had no major impact on the high-temperature-induced repression of lateral root development (Supplementary Fig. 22a-e). We accordingly conclude that *PHYB* and *CRY1* jointly mediate the light-modulated integration of ambient temperature into the root system architecture.

*PHYB* and *CRY1* both mediate light signals by regulating *PHYTOCHROME INTERACTING FACTORS* (*PIFs*) and *ELONGATED HYPOCOTYL5* (*HY5*) module, which is also known to integrate light quality during the shade avoidance response into the rate of lateral rooting^26,27,28^. However, neither the quadruple mutant *pif1,3,4,5* (*pifQ*), nor *hy5-215* mutant was distinguishable from the wild type in regards to HT-dependent lateral rooting under ML condition (Supplementary Fig. 22a and f-i). This suggests a molecularly distinct mechanism for the integration of light quality during shade avoidance and the light quantity-dependent modulation of HT into lateral rooting.

### Concluding remarks

Mechanistically, we propose that the PHYB- and CRY1-mediated light quantity in the shoot defines the strength of a systemic oscillatory auxin signal in the main root, thereby defining the auxin-dependent identity of lateral root PBS. The light-dependent, systemic auxin dynamics also modulate HT-dependent control of PBS progression to lateral roots, suggesting a general mechanism for environmental signal integration. However, the mechanism by which shoot-sensed light signalling modulates systemic oscillatory auxin signals in the root remains to be addressed. Light receptors are well known for shaping plant architecture by modulating key components of the circadian clock^29^. Notably, the 24-hour rhythm of systemic auxin oscillations may pinpoint a potential role for the circadian clock in signal transmission. While this aspect requires further investigation, it is noteworthy that the circadian clock regulates at least some auxin-mediated transcriptional responses^30^. Conceptually, our work illustrates that two oscillatory systems define LR spacing. While a local auxin oscillation zone regulates the regular priming of PBS during main root growth, the systemic oscillatory auxin signal along the main root integrates environmental information, such as light and temperature, to control the PBS identity in time and eventually its progression to become lateral roots. This framework reveals how plants mechanistically use distinct oscillatory signals of the same regulator to combine robustness and plasticity into *de novo* organogenesis.

## Materials and methods

### Plant Materials

The *Arabidopsis thaliana* ecotype *Columbia 0* (*Col-0*) and *Landsberg* (*Ler*) were used as the wild type in this study. The *pDR5::Luc*^2^ and *slr-1*^21^ are in the ecotype *Col-0* accession and was received from Stefan Kircher. The mutants of *phyA-211*^31^, *phyB-9*^32^, *cry1-304*^33^, *pif1,3,4,5*^34^ and *hy5-215*^35^ mutants are all in the *Col-0* accession and were received from Andreas Hiltbrunner. The mutant of *phyBcry1cry2*^36^ is in the ecotype Ler accession and was received from Stefan Kircher. *pIAA14::Luc* and *p35S::Luc* were created using floral dipping in the *Col-0* background. *slr-1×pDR5::Luc* was generated by crossing *slr-1* with *pDR5::Luc*.

### Plasmid construction and plant transformation

To construct the *p35S::Luc* control plasmids, the *CaMV35S* promoter was inserted upstream of the *Luciferase* gene in the pPLV03 vector^37^. Genomic DNA from Arabidopsis ecotype Col-0 was used as the template for amplification of the upstream promoter of AT4G14550 (IAA14) using the primers forward 5’-ACAACTTTGTATAGAAAAGTTGTATGCCTCTCCACTACCACT −3’ and reverse 5’-ACTGCTTTTTTGTACAAACTTGCTCTTCTTGCTGTCTATATATGT −3’ and the PCR product cloned into the pDONR P4-P1 vector (Invitrogen) and subsequently recombined with the pENTR/D-TOPO-Luciferase clone^2^ into the dpGreen-BarT destination vector^38^. All constructs were verified by sequencing and introduced into *Agrobacterium tumefaciens* strain GV3103 by electro-transformation, and subsequently transformed into the *Col-0* ecotype background by floral dipping. The harvested seeds were sown on soil and sprayed 0.2g/L Basta herbicide solution (Bayer) for selecting positive *p35S::Luc* or *pIAA14::Luc* seedlings.

### Growth conditions and treatments

The Arabidopsis seeds were sterilized for 2–5 minutes with 70% ethanol, followed by drying. After sterilization, the seeds were uniformly plated on one single line on square plates (12 × 12 × 1.5 cm). The plates contained 50 mL standard Murashige and Skoog solid medium, which is made of 0.8% agar (Duchefa), 0.5× Murashige and Skoog (MS) medium (Duchefa), and with 1% sucrose (MS+, no exogenous sucrose was added for MS-) (pH 5.9). Subsequently, the seeds were stratified for 2 days in 10 °C and dark conditions. Seeds were germinated on vertically positioned plates within a Weiss-Technik incubator (Fitotron SGC 2), exposed to top-providing illumination from LED white cultivation lights (16h day/8h night cycle) in 21 °C and an irradiance of 120 µmol m^−2^ s^−1^. Plates grow under the above control condition for 4 days (root quantification assay) or 5 days (luminescence monitoring assay) before subsequent experiments. For light conditions and HT treatment, two Weiss-Technik incubators were configured at either 21°C (control) or 29°C (HT), operating under constant light. Each incubator was outfitted with top LED white culture lights arranged across three distinct culture levels, featuring irradiances of 20, 50, and 120 (control) µmol m^−2^ s^-1^.

### Root quantification

For root phenotyping, surface-sterilized seeds were uniformly plated, stratified, and germinated on a square plate (12 × 12 × 1.5 cm) as described above. After germination under the control condition for 4 days, marking the position of the root tip, the 4-day-old seedlings were then transferred to the treatment condition (either 21°C or 29°C with light conditions set at 20, 50 or 120 µmol m^−2^ s^-1^) for additional 3 days and the root tip positions were marked again. For IAA and Auxinole treatment, water solution containing 0.2 μM IAA (OlChemIm s.r.o.) or 20 μM Auxinole (OlChemIm s.r.o.) was evenly spread on the plates, with solvent (DMSO, Duchefa) used as a control. For lateral and primary root phenotyping, the plates were scanned at 600 dpi using an Epson V850Pro photonegative scanner at 24-hour intervals, starting from the 3rd day to the 10th day of transferring. The lateral roots between the two marked root section were counted manually from the images and the length of primary roots between two marked positions was measured automatically using the SNT plugin or manually with Fiji/ImageJ. The significance test was conducted in R using the t.test and aov functions.

### Lateral root primordia quantification

To phenotype lateral root primordia, seeds were placed on individual square plates. After marking the root tips of 4-day-old seedlings, they were moved to specific treatment conditions as outlined in the manuscript and figure legend (either 21°C or 29°C with light conditions set at 50 or 120 µmol m^−2^ s^-1^). Following a 3-day treatment, the positions of the root tips were marked again, and the plates were scanned to measure the length of the primary roots. After 5 days of treatment, primary roots located between two marked positions were harvested, followed by fixation and clearing in accordance with a previously published protocol^39^. The roots were mounted in 50% glycerol and analyzed using a Zeiss AxioObserver Z1 microscope (Plan-Apochromat 40×/0.95 objective) with a AxioCam MRc camera. Primordia were counted manually, and the length of primary roots between two marked positions was determined as described above. The significance test was conducted in R using the aov functions.

### Luminescence imaging and analysis

The luciferase activity in roots was visualized by time-lapse imaging of seedlings sprayed with 2 mM luciferin (Biosynth AG, Switzerland). *DR5::Luciferase* seedlings were grown using the general procedure as described above. To perform time-lapse imaging of the *DR5:Luciferase* expression in the oscillation zone, a Vers Array XP camera system (Roper Scientific) was used to image the luciferase signal in the vertical growing Arabidopsis root tip from 5 days old (4 to 8 seedlings) with the set-up of the exposure times of 5 min (binning 2) interrupted by 14 min LED WL (measured irradiances of 10∼30 μmol·m^-2^·s^-1^ for low light, 40∼60 μmol·m^-2^·s^-1^ for medium light and 110∼130 μmol·m^-2^·s^-1^ for high light condition) followed by 1 min darkness. The temperature in the imaging chamber was configured at either 21°C or 29°C by a recirculating Cooler (JULABO FL300). Image sequences were saved for further analysis in Fiji/ImageJ. Image sequences were converted into kymographs by tracking the final course of root growth. This visualization captures spatiotemporal changes in Luciferase signals during primary root growth. To quantify the universal *DR5* signal in primary roots, the grey value of the primary root excluding the tip to the oscillation zone was recorded from the kymograph at an interval of every 10 hours. The PBS were counted manually from the kymographs and original image sequences. The length of primary roots was measured manually from the original images. To determine the time and strength of the systemic *DR5* signal in primary roots, the time point and grey value were recorded when the systemic *DR5* signal in the primary roots reached the top value. To imaging luminescence of *p35S::Luc* control lines, seedlings were grown to 5-days-old on solid MS+ medium sprayed with 2 mM luciferin (Biosynth AG, Switzerland) following the general procedure as described above. Seedlings were then transferred into Weiss-Technik incubators set to either 21°C (control) or 29°C (HT), with constant illumination of 120 μmol·m^-2^·s^-1^ as described above. The images of luminescence in primary root were captured using a FUSION SL4 system (VILBER), with 30 second exposure at corresponding time points. The gray value of the primary root, excluding the tip up to the oscillation zone (10 mm from root tip), was measured to assess the universal luminescence signal in primary roots. All measurements were performed manually with Fiji/ImageJ. The significance test was conducted in R using the t.test and aov functions. The Anderson-Darling normality test was conducted in R using the ad.test function.

## Acknowledgements

We thank Stefan Kircher and Andreas Hiltbrunner for providing seed materials and critically reviewing the manuscript; Stefan Kircher for assistance with root luminescence imaging and analysis; Deutsche Forschungsgemeinschaft (DFG) (DFG; 470007283 to J.K.-V. and CIBSS – EXC-2189; 390939984 to J.K.-V.), Chinese Scholarship Council (CSC, 202006300036 to C.R.) and MICIU/AEI/10.13039/501100011033 (CEX2020-000999-S to M.A.M.-R. through CBGP) for funding.

## Author contributions

C.R. and J.K.-V. designed the research; C.R. and J.B. performed the research; C.R. analyzed data; C. R. generated the *p35S::Luc* plasmid and plant material and M.A.M.-R. generated the *pIAA14:Luc* and *slr-1 × pDR5::Luc* plasmids and plant material; and C.R. and J.K.-V. wrote the paper. All authors edited the paper.

---

**Supplementary Figure 1.**
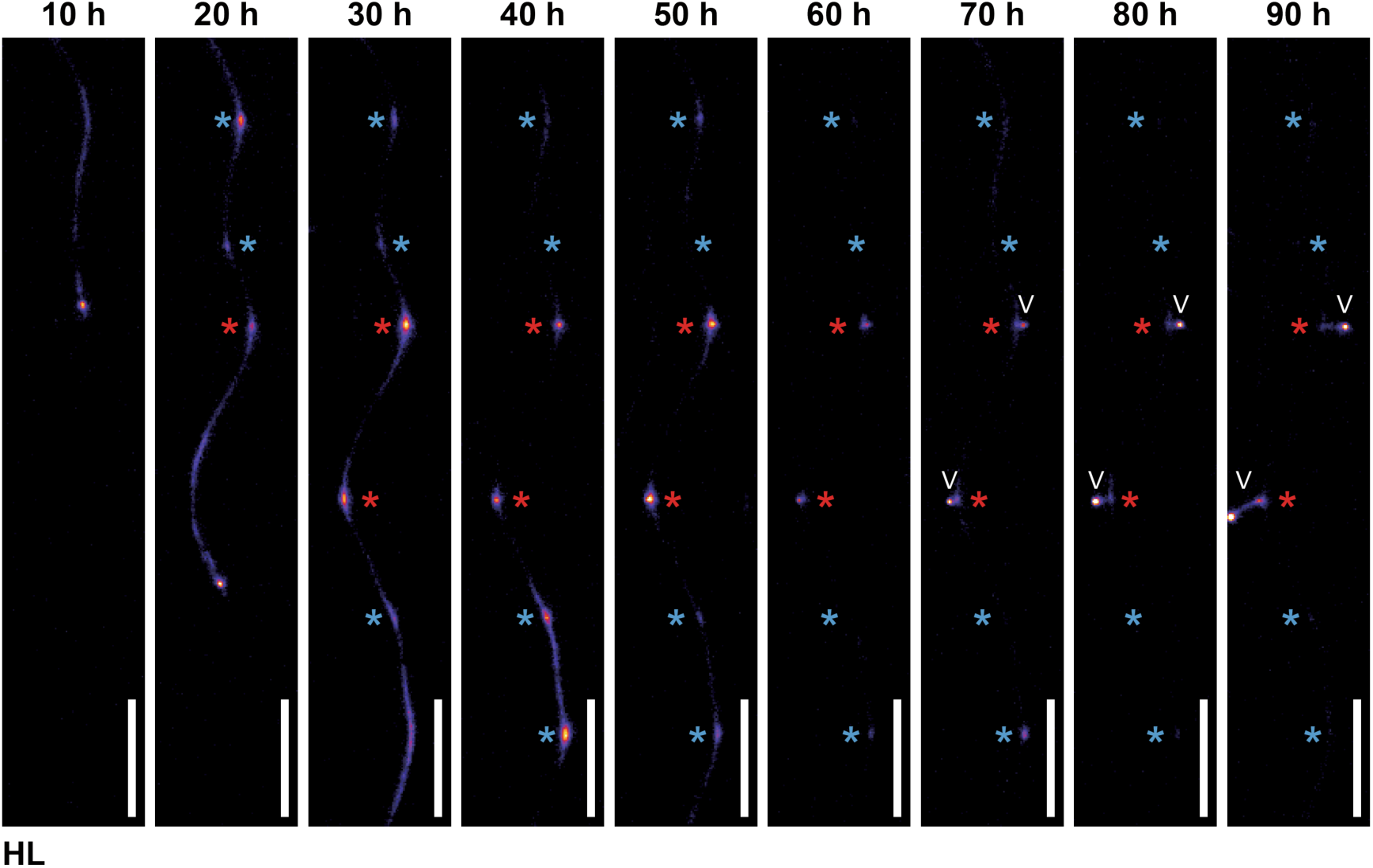
Original images showing the distinct progression of persistent and transient PBS at HL condition. Time series images of the primary root were taken every 10 hours under 21°C and high light (HL) conditions. Persistent PBS is marked by a red asterisk and transient PBS is marked by a blue asterisk. The white arrowhead denotes the emerged lateral root (Scale bar = 1 mm).

**Supplementary Figure 2.**
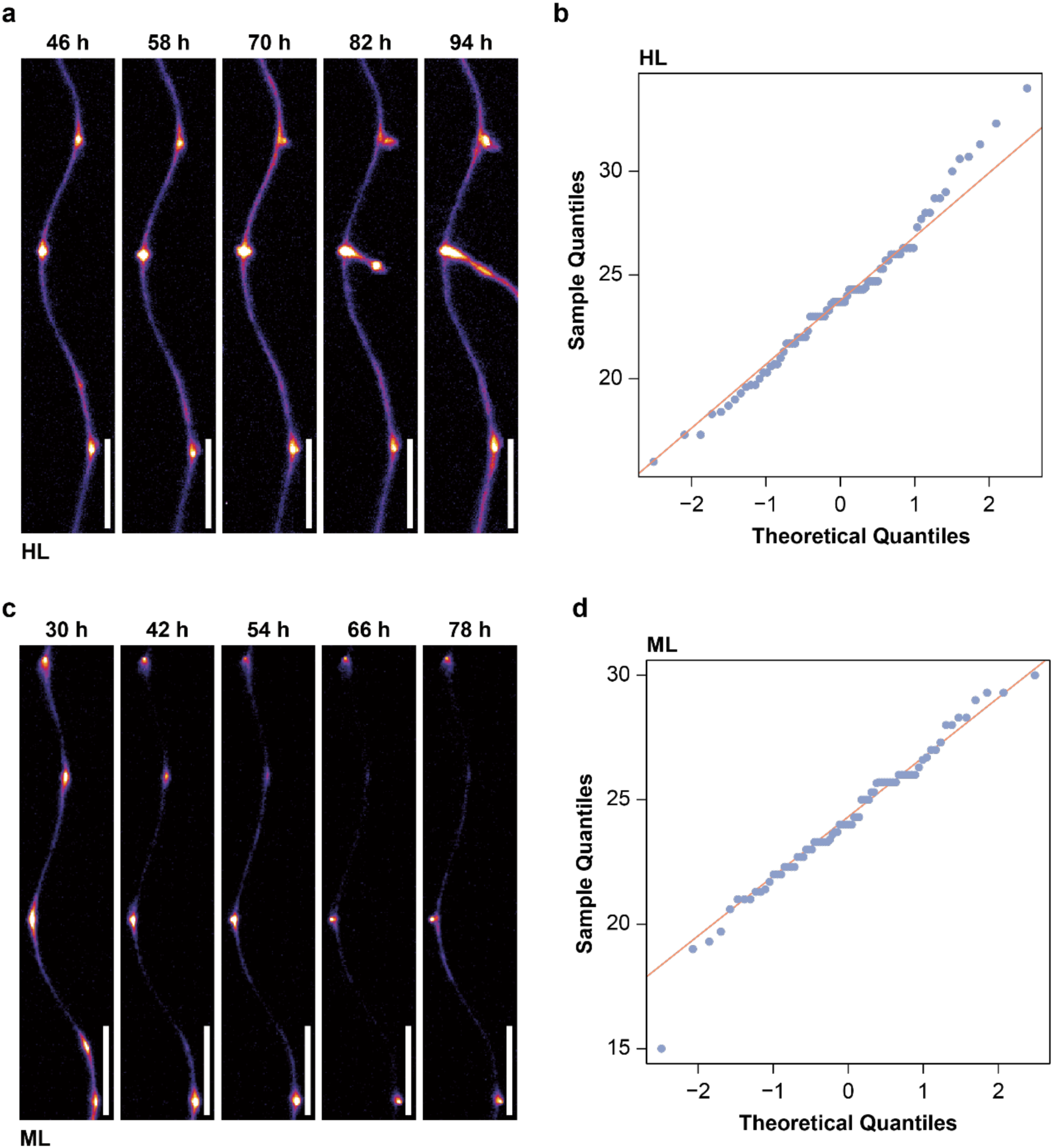
The time intervals between oscillatory peaks of systemic auxin follow a normal distribution. **a**, **c**, Time series images of the primary root were taken every 12 hours under high light (HL) (**a**) and medium light (ML) (**c**) at 21°C conditions (Scale bar = 1 mm). **b**, **d**, The quantile-quantile (Q-Q) plot compares the sample quantiles of the time intervals between peaks of systemic auxin oscillations with the corresponding theoretical quantiles of normal distribution at 21°C and light intensities of HL (**b**) and ML (**d**) conditions (*n* = 23 to 29). The pattern closely following the orange line indicates that the frequency of the systemic oscillation is likely to follow a normal distribution.

**Supplementary Figure 3.**
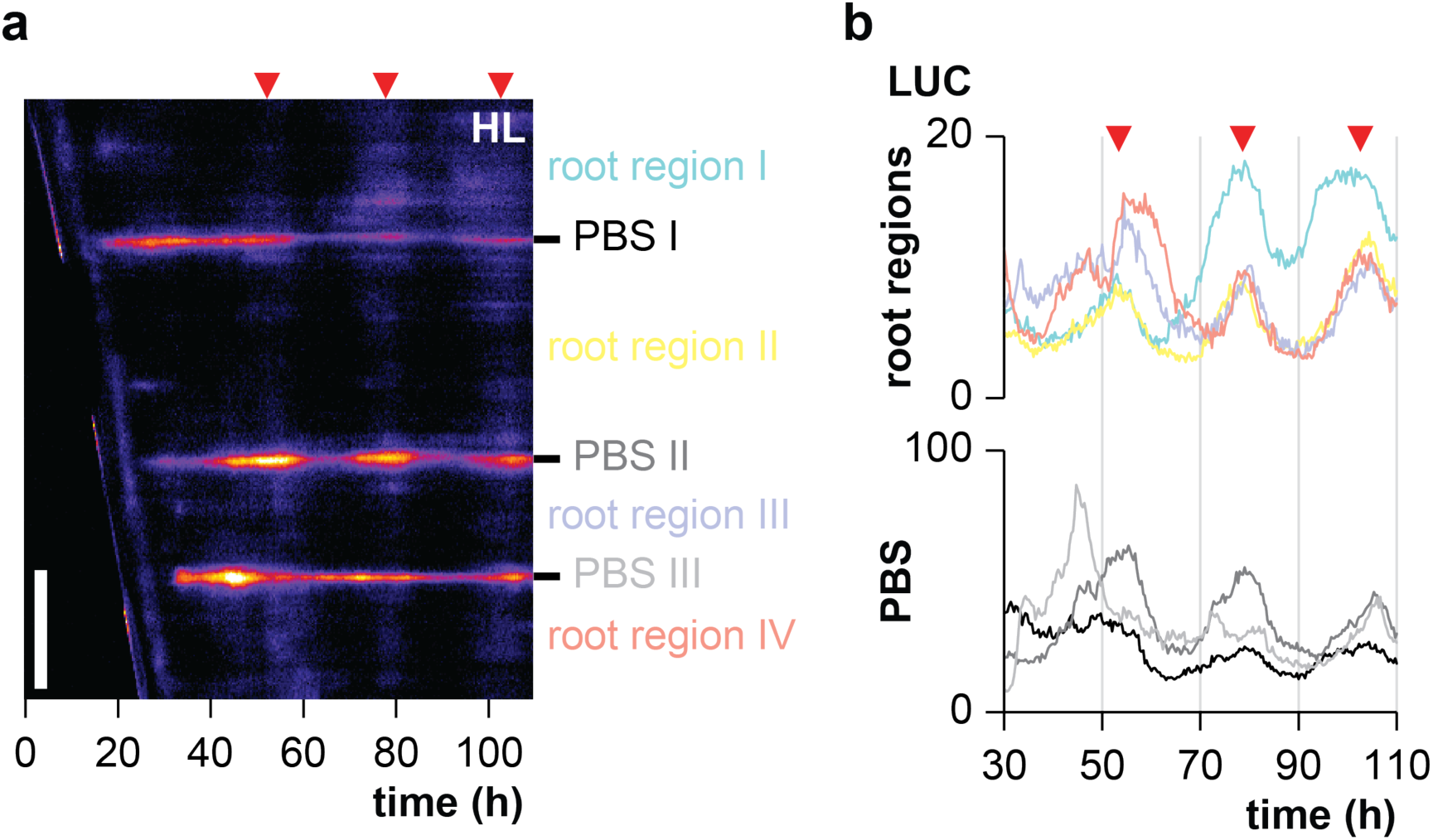
Kymograph showing the distinct progression of persistent and transient PBS at HL condition. **a**, **b**, Kymograph (**a**) and real-time quantification (**b**) of luminescence display auxin signalling dynamics in specific root regions and PBS, with red triangles mark oscillatory peaks of systemic auxin signal (Scale bar = 1 mm). The colored notation on the right of (**a**) marks the quantified root regions and PBS, corresponding to the same colors in (**b**).

**Supplementary Figure 4.**
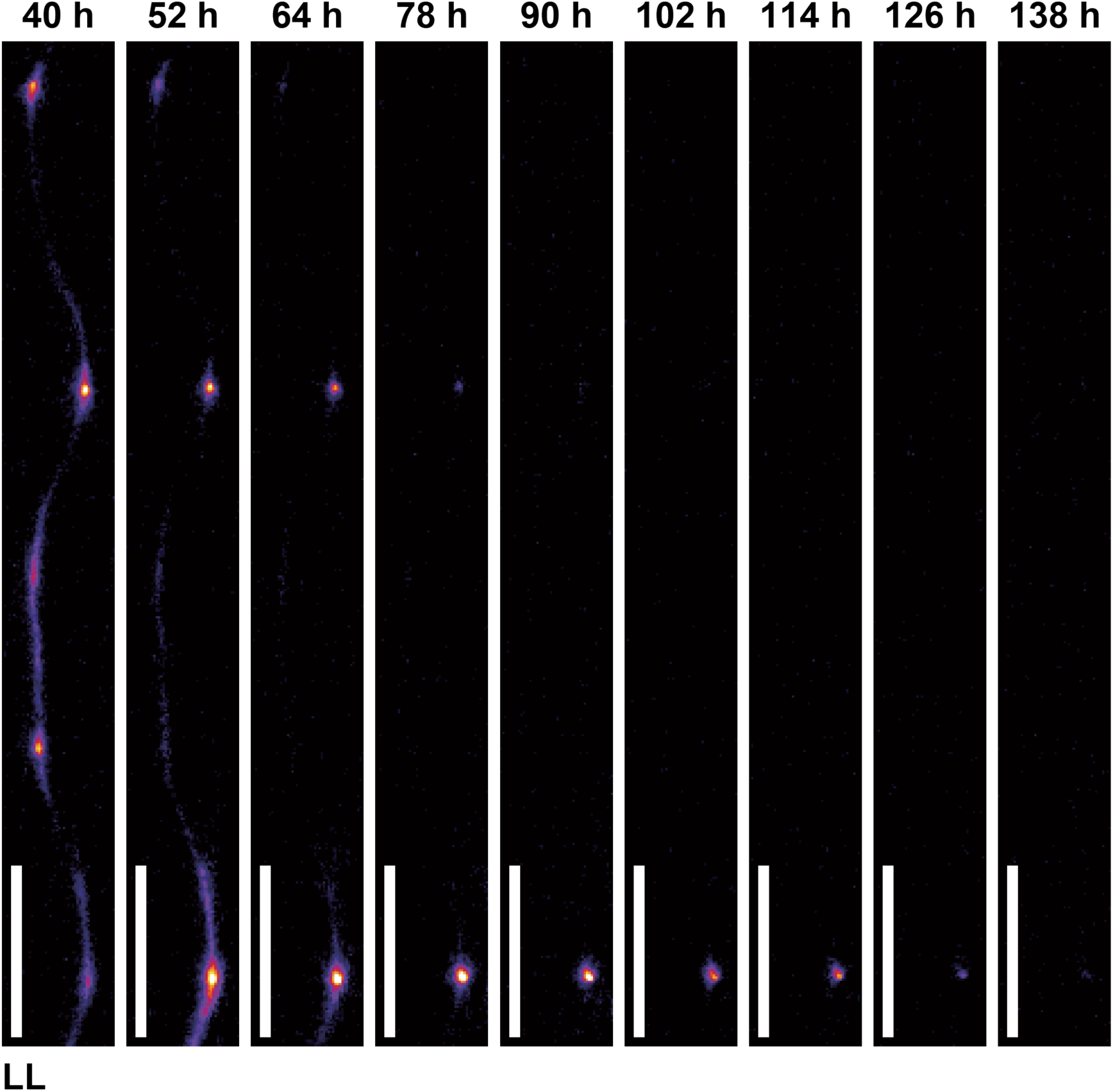
Original images showing the dynamic of systemic auxin signal and progression of PBS at low light conditions. Time series images of the primary root were taken every 12 hours under 21°C and low light (LL) conditions (Scale bar = 1 mm).

**Supplementary Figure 5.**
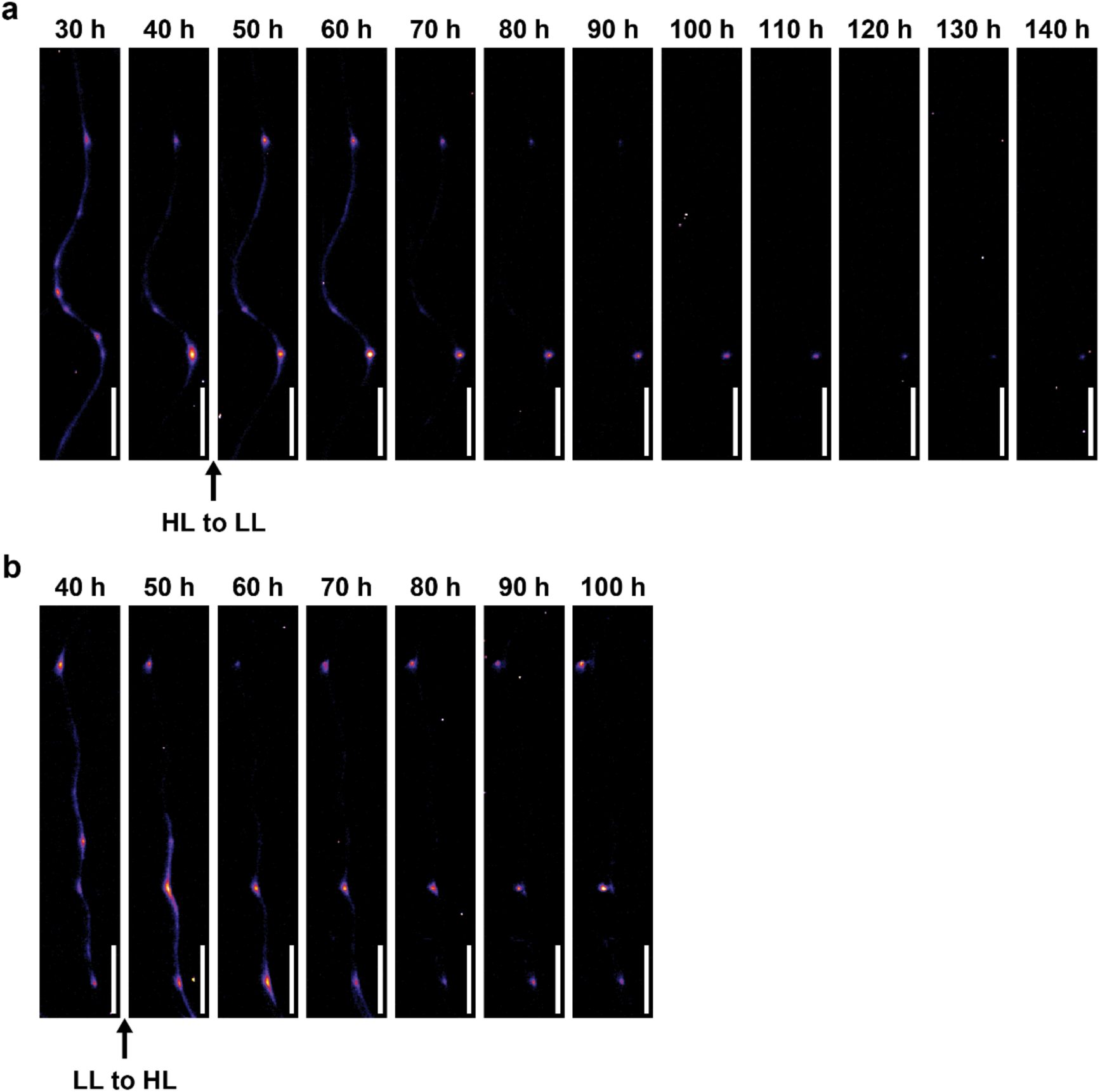
Original images showing the dynamic of systemic auxin signal and progression of PBS during light transitions. **a**, **b**, Time series images of the primary root were taken every 12 hours under light transitional conditions from high light (HL) to low light (LL) (**a**), and from LL-to-HL (**b**) (Scale bar = 1 mm).

**Supplementary Figure 6.**
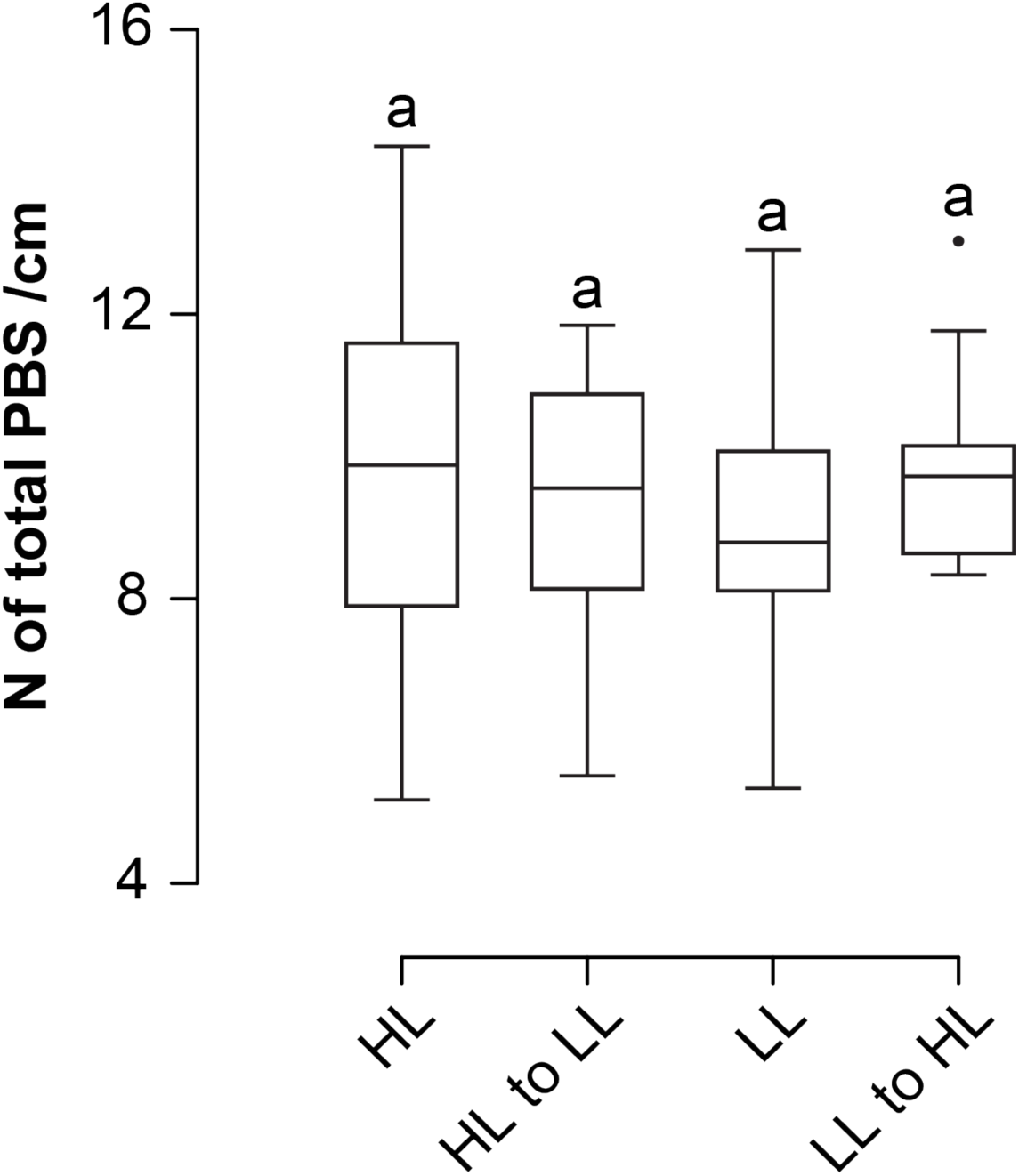
The density of total PBS is quantitatively comparable at maintained and fluctuating light conditions. Quantification of total PBS density under high light (HL), low light (LL) and transfer from HL-to-LL and LL-to-HL conditions (*n* = 16 to 38). Letters indicate values with statistically significant differences from one-way ANOVA performed (*P* > 0.05).

**Supplementary Figure 7.**
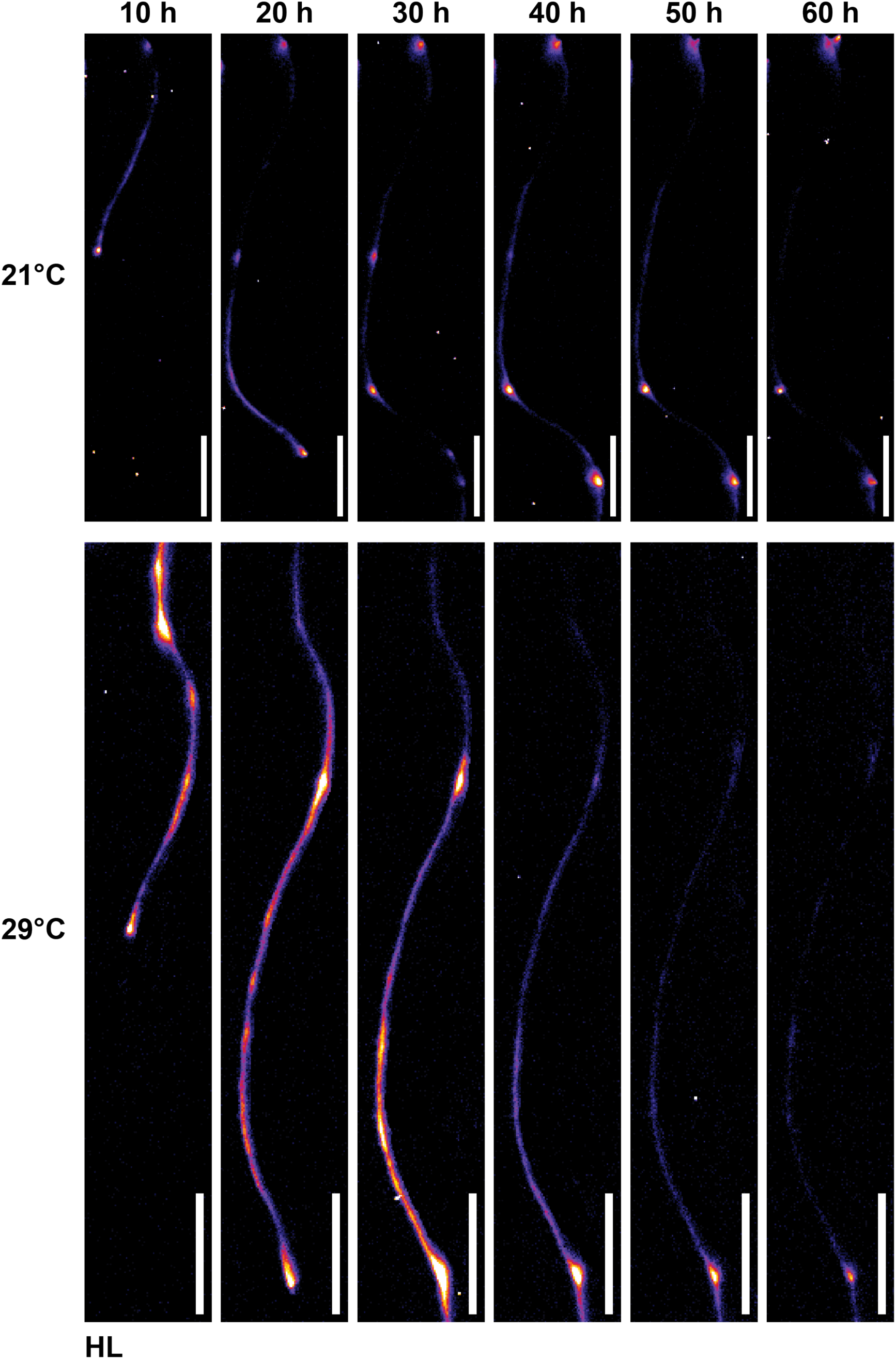
Original images showing the dynamic of systemic auxin signal and progression of PBS under 21°C and 29°C at high light conditions. Time series images of the primary root were captured every 10 hours under high light (HL) conditions at 21°C (top row) and 29°C (bottom row) (Scale bar = 1 mm).

**Supplementary Figure 8.**
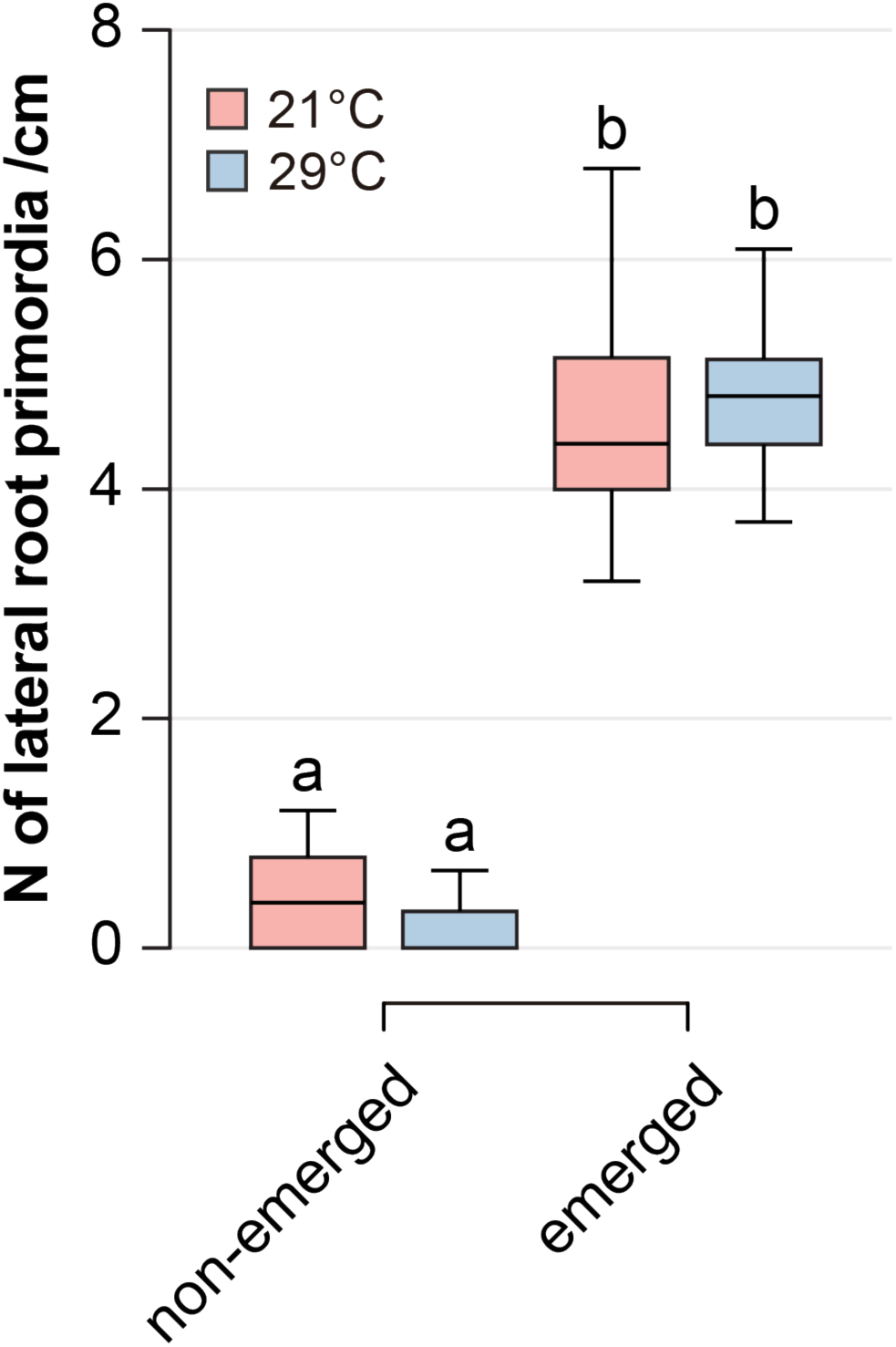
Density of lateral root primordia is quantitively comparable between 21°C and 29°C at high light conditions. Quantification of lateral root primordia density at 21°C and 29°C under high light conditions (*n* = 34 to 38). Letters indicate values with statistically significant differences from one-way ANOVA performed (*P <* 0.0001).

**Supplementary Figure 9.**
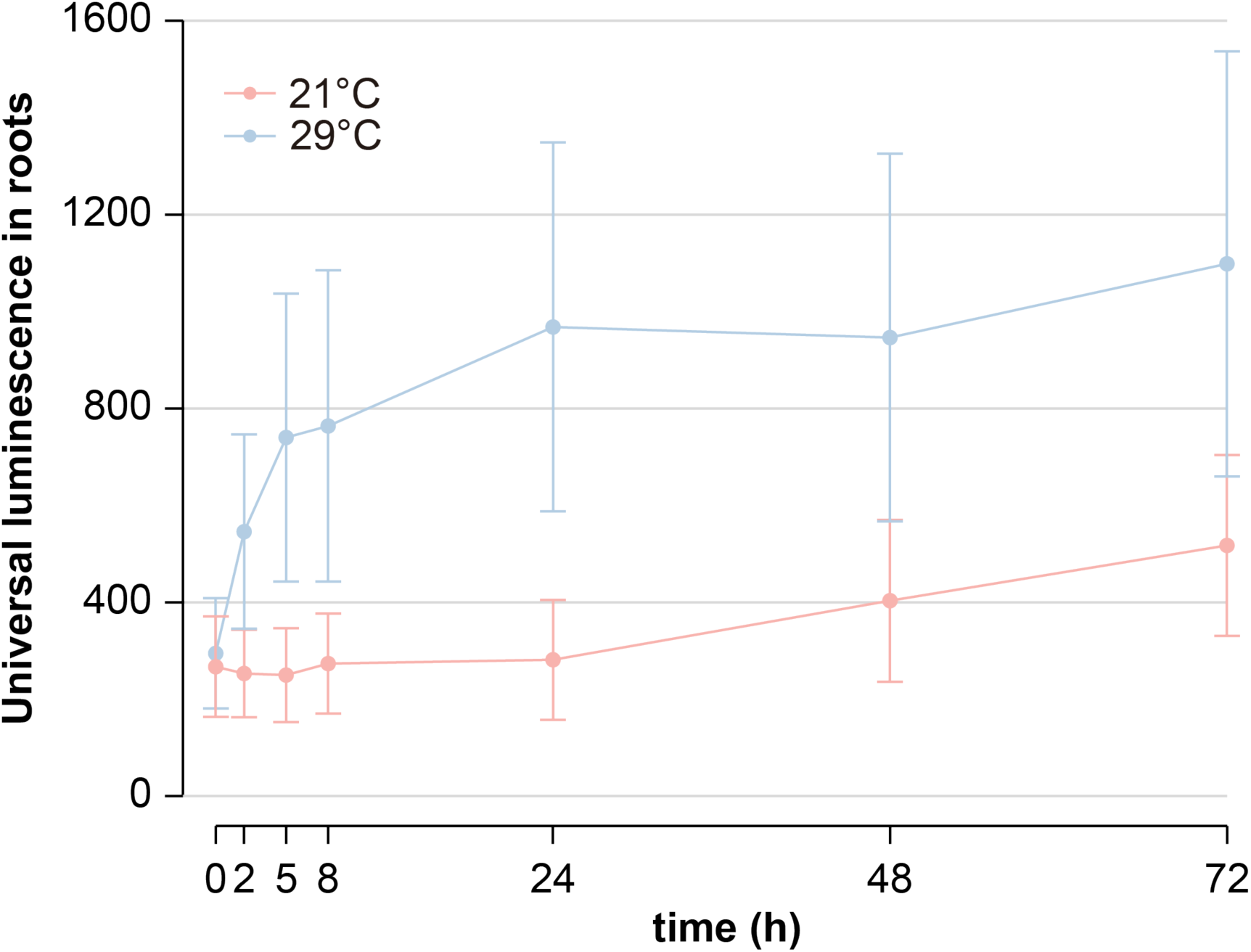
Time series quantification of universal *p35S::Luc* signal in primary roots at 21°C and 29°C. Quantification of luminescence in primary roots showing the dynamics of the *p35S::Luc* signal at 21°C and 29°C under high light conditions (Error bars represent standard deviation; *n* = 45 to 47).

**Supplementary Figure 10.**
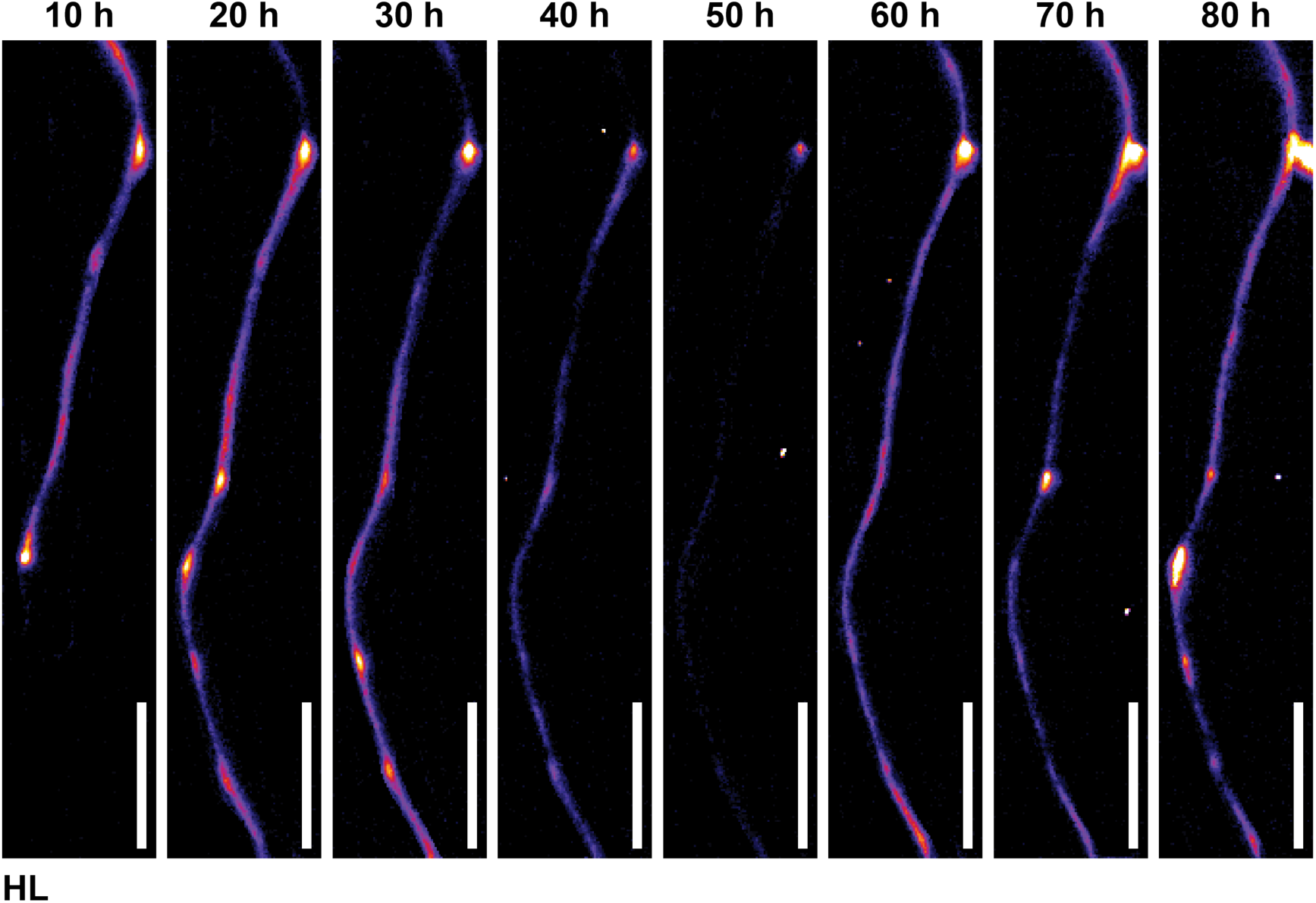
Original images showing the recovery of systemic auxin signal and PBS under high temperature at high light conditions. Time series images of the primary root were captured every 10 hours under high light (HL) conditions at 29°C. Scale bar = 1 mm.

**Supplementary Figure 11.**
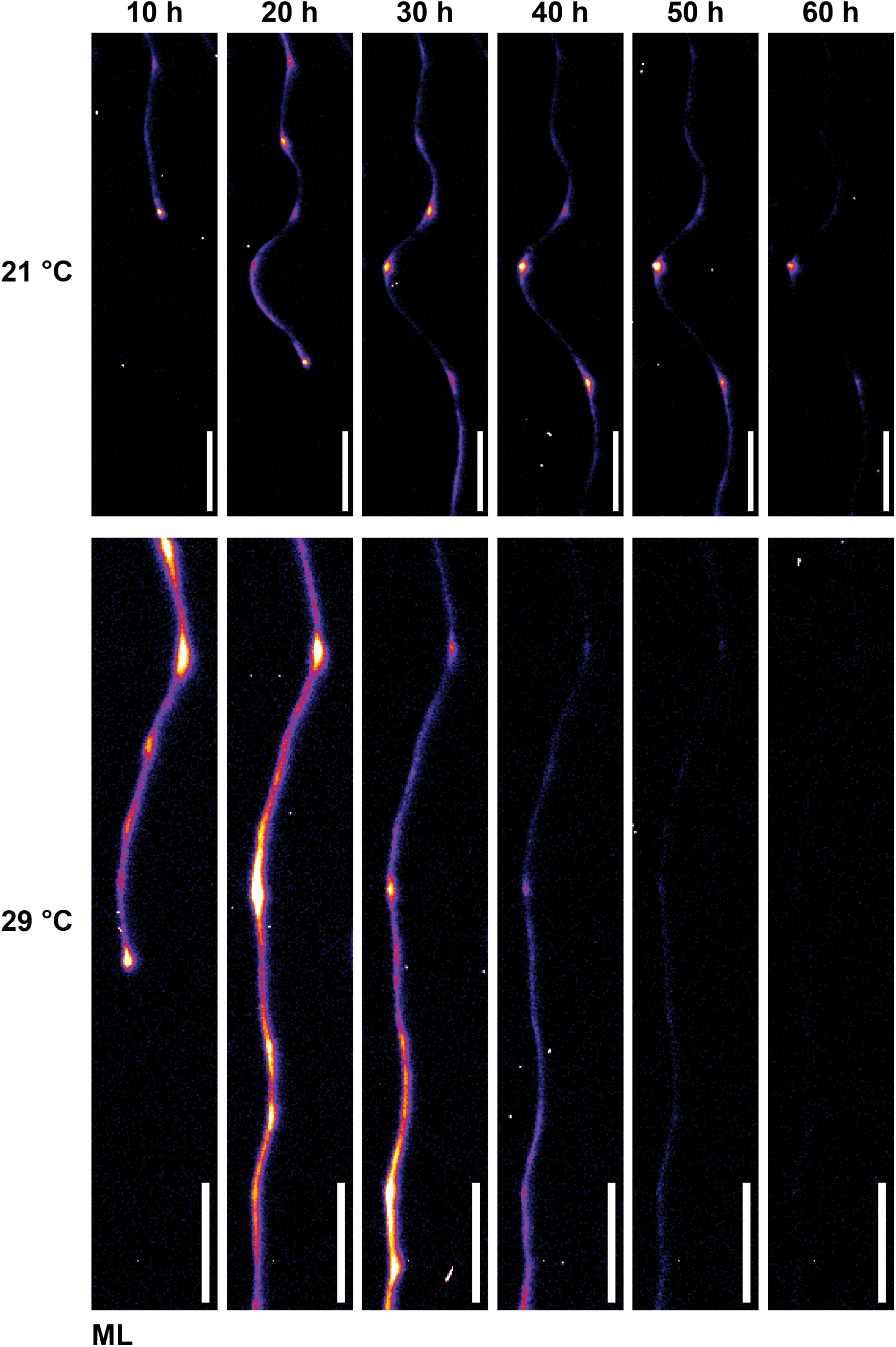
Original images showing the dynamic of systemic auxin signal and progression of PBS under 21°C and 29°C at medium light conditions. Time series images of the primary root were captured every 10 hours at 21°C (top row) and 29°C (bottom row) under medium light (ML) conditions (Scale bar = 1 mm).

**Supplementary Figure 12.**
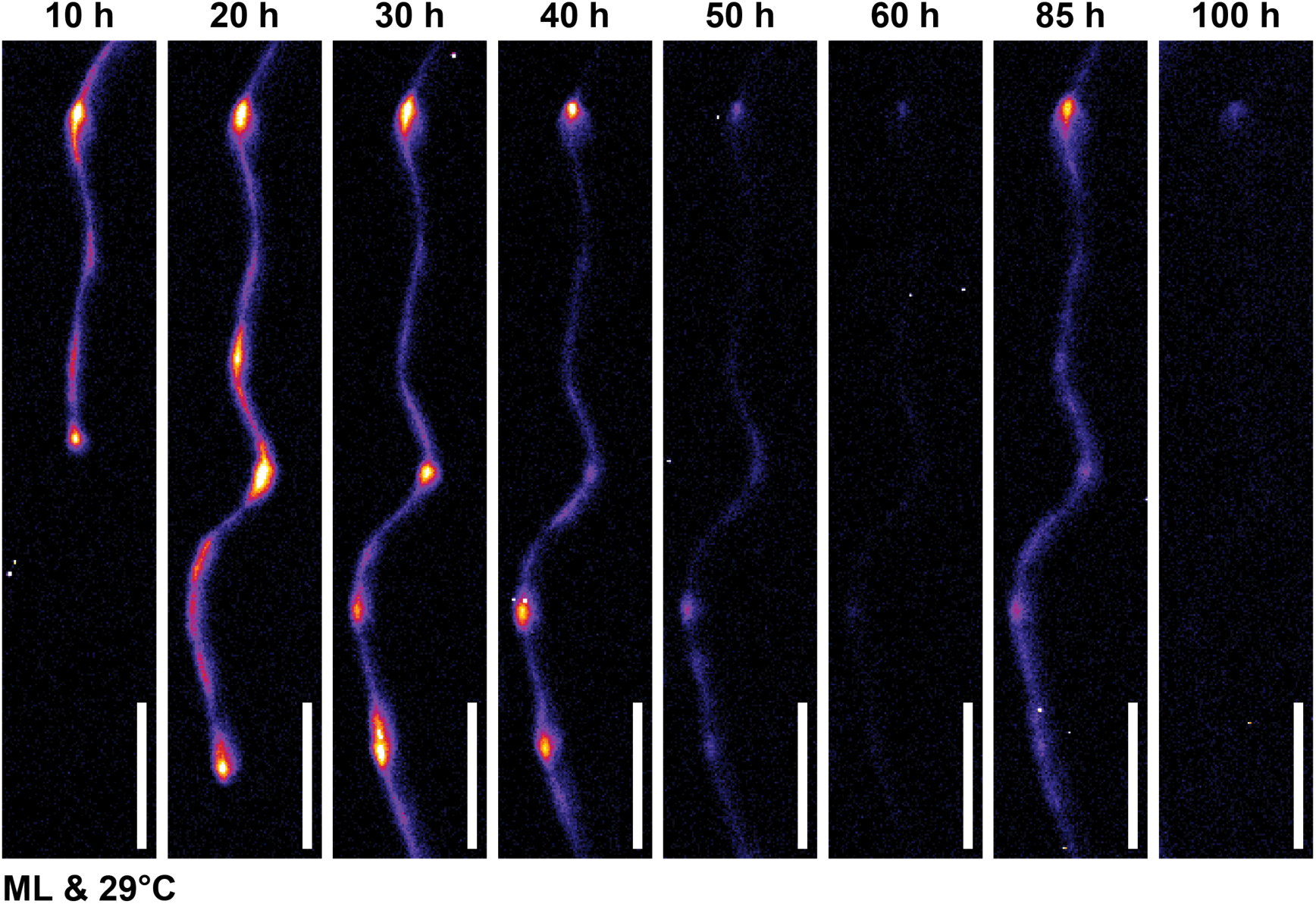
Original images showing the recovery of systemic auxin signal and PBS under high temperature at medium light conditions. Time series images of the primary root were captured every 10 hours under 29°C and medium light (ML) conditions (Scale bar = 1 mm).

**Supplementary Figure 13.**
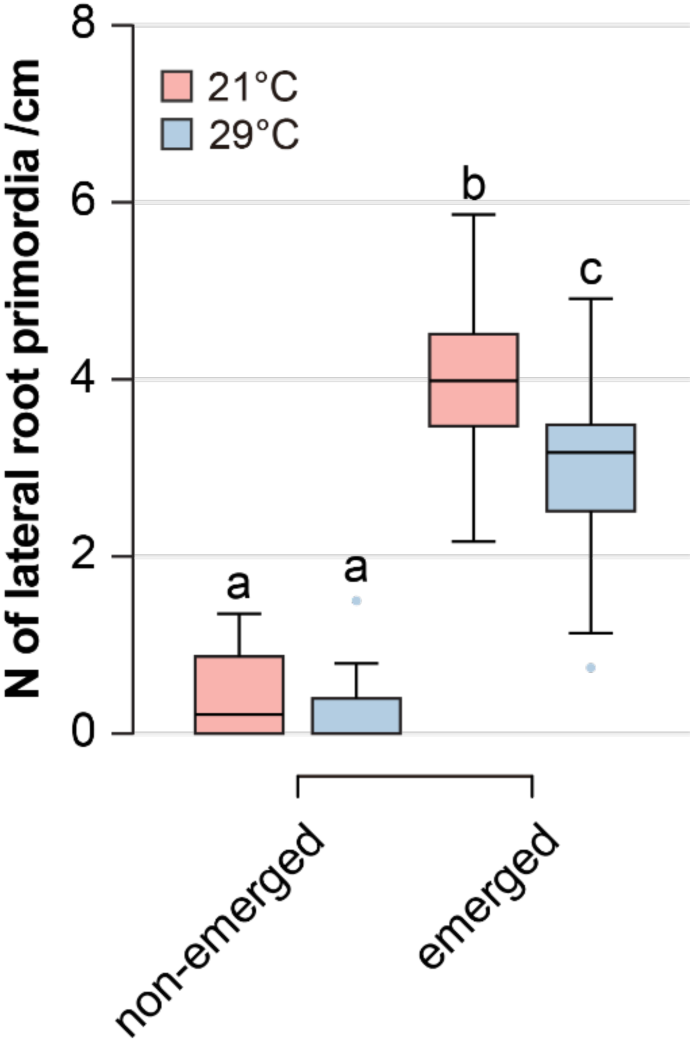
Density of lateral root primordia under 21°C and 29°C at medium light conditions. Quantification of lateral root primordia density at 21°C and 29°C under medium light conditions (*n* = 32 to 34). Letters indicate values with statistically significant differences from one-way ANOVA performed (*P <* 0.0001).

**Supplementary Figure 14.**
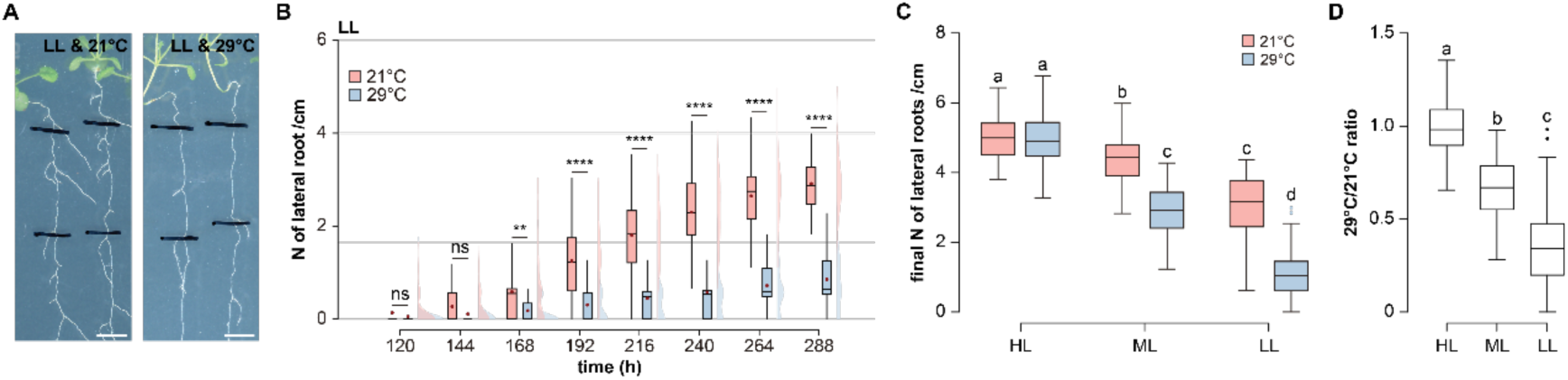
Light quantity modulates the high temperature-dependent inhibition of lateral root density. **a**, **b**, Representative images of root systems (**a**) and time series quantification of LR density (**b**) at 21°C and 29°C under low light (LL) conditions (*n* = 53 to 55; Scale bar = 5 mm). **c**, **d**, Quantification (**c**) and calculated ratio (**d**) of final LR density at 21°C and 29°C under high light (HL), medium light (ML) and LL conditions (*n* = 53 to 55). Letters indicate values with statistically significant differences from one-way ANOVA performed for (**c**) and (**d**) (*P <* 0.0001 for both).

**Supplementary Figure 15.**
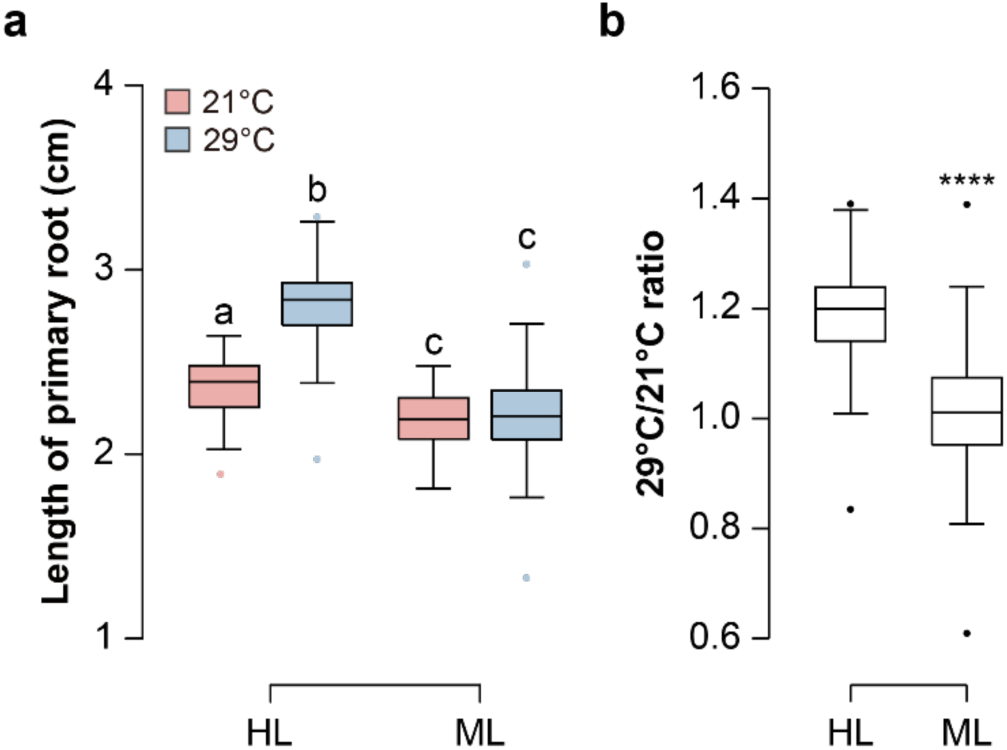
High temperature-dependent elongation of the primary root relies on light. **a**, **b**, Quantification (**a**) and Calculated ratio (**b**) of primary root length of *Col-0* at 21°C and 29°C under high light (HL) and medium light (ML) (*n* = 54 to 55). Letters indicate values with statistically significant differences from one-way ANOVA performed for (**a**) (*P <* 0.0001). Paired and two-tailed student’s t-test performed for (**b**) (*P <* 0.0001****).

**Supplementary Figure 16.**
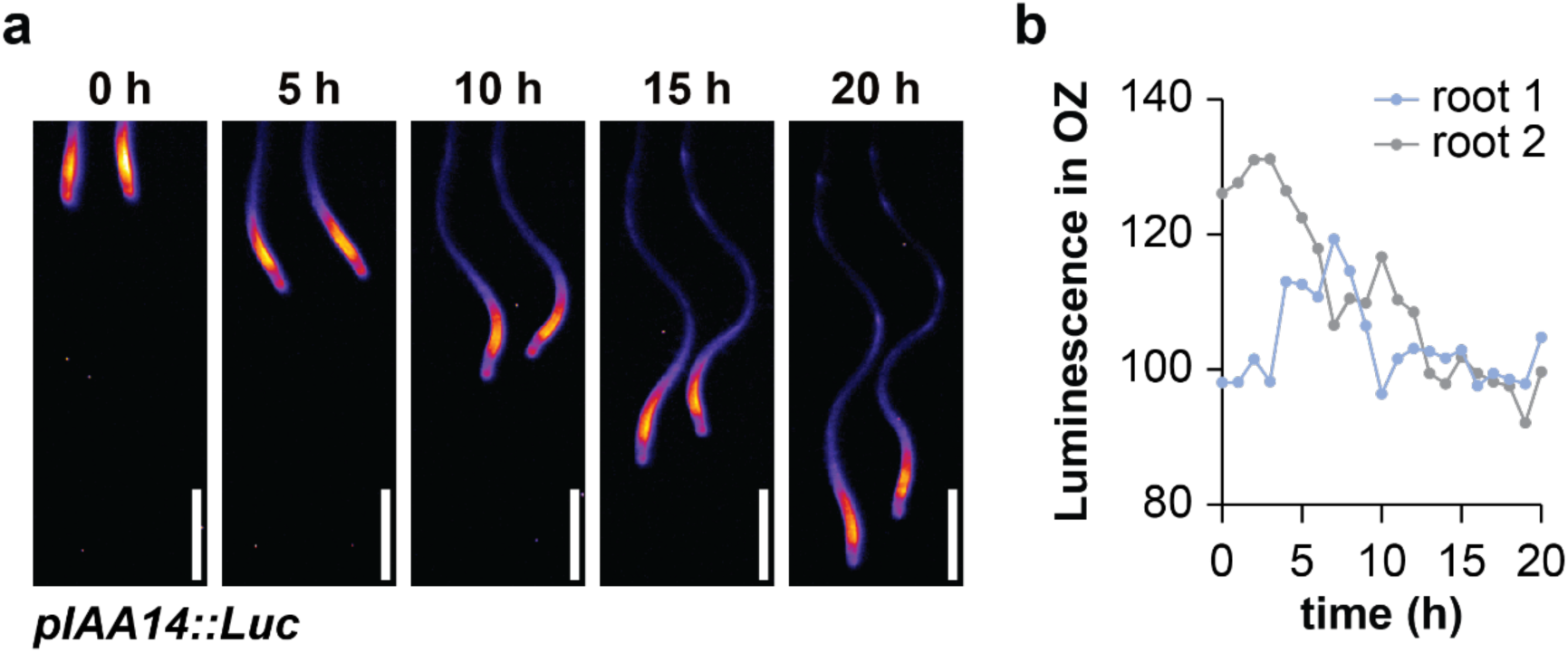
Original images and quantification showing the dynamic of *IAA14* reporter in oscillation zone. **a**, Time series images of the primary root were captured every 5 hours under 21°C and HL conditions (Scale bar = 1 mm). **b**, Time series luminescence quantification in the oscillation zone (OZ) of two independent roots.

**Supplementary Figure 17.**
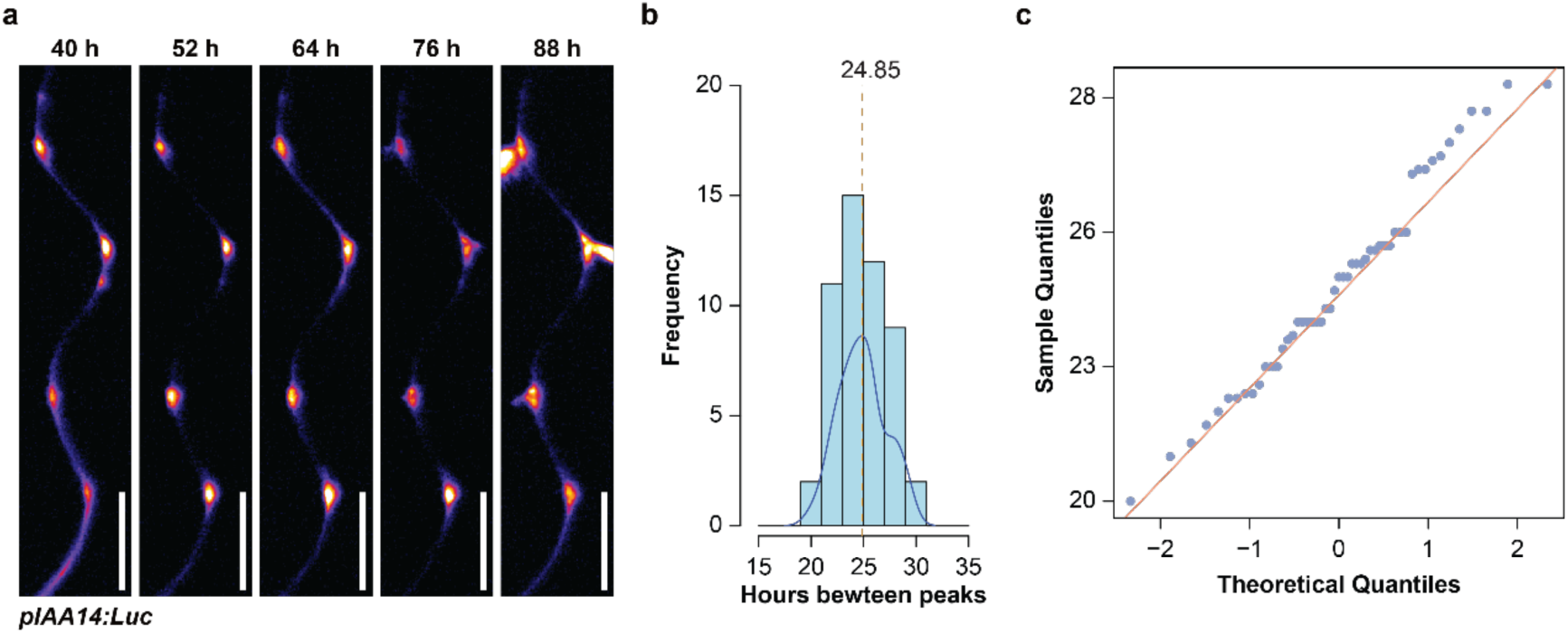
The time intervals between oscillatory peaks of *IAA14* reporter signal follows a normal distribution. **a**, Time series images of the *pIAA14::Luc* primary root were taken every 12 hours under high light (HL) at 21°C conditions (Scale bar = 1 mm). **b**, Quantified distribution of the time intervals between oscillatory peaks of *IAA14* reporter signal under high light (HL) at 21°C conditions (mean value marked by a yellow dashed line; *n* = 17). **c**, The quantile-quantile (Q-Q) plot compares the sample quantiles of the time intervals between peaks of *IAA14* reporter oscillations with the corresponding theoretical quantiles of normal distribution at 21°C and light intensities of HL (*n* = 17). The pattern closely following the orange line indicates that the frequency of the systemic oscillation is likely to follow a normal distribution (Anderson-Darling normality test, P-value = 0.4336).

**Supplementary Fig. 18.**
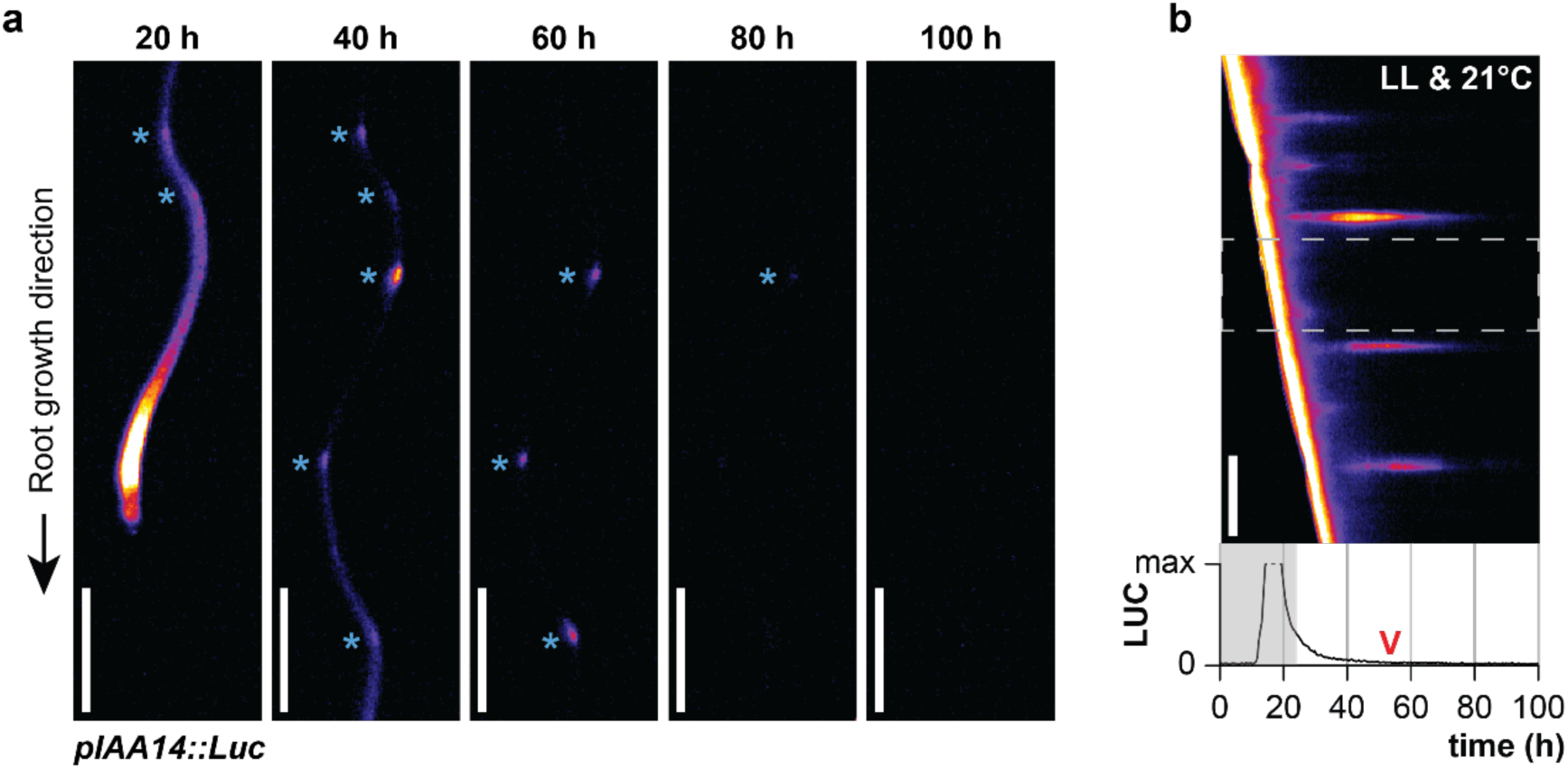
Original images and kymograph showing the dynamic of *IAA14* reporter signal and progression of PBS at low light conditions. **a**, Time series images of the *pIAA14::Luc* primary root under low light (LL) at 21°C conditions (Scale bar = 1 mm). Blue asterisks mark transient PBS. **b**, Kymograph showing dynamic of *pIAA14::Luc* and progression of PBS under LL at 21°C conditions (Scale bar = 1 mm). Quantification of real-time luminescence is below the kymographs where the red arrowhead denotes the downregulation of the *pIAA14::Luc* signal. The dashed grey rectangle in the kymographs mark the region for the real-time plots. The shaded area in real-time plots marks the signal originating from the root tip and local oscillation (root clock) zone.

**Supplementary Fig. 19.**
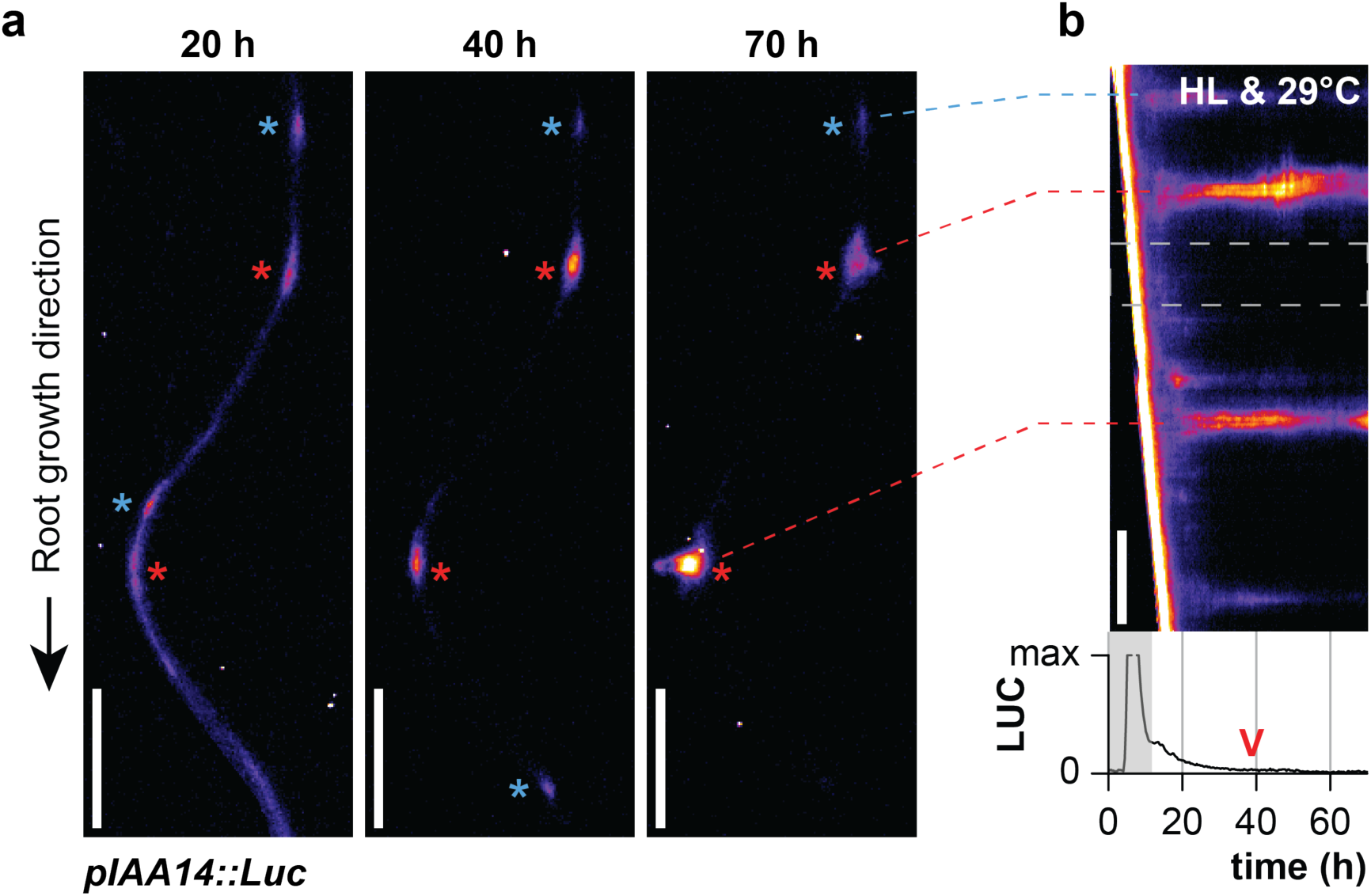
Original images and kymograph showing the dynamic of *IAA14* reporter signal and PBS under high temperature and high light conditions. **a**, Time series images of the *pIAA14::Luc* primary root under high light (HL) at 29°C conditions (Scale bar = 1 mm). Red asterisks mark persistent PBS and blue asterisks mark transient PBS. **b**, Kymograph showing dynamic of *pIAA14::Luc* and progression of PBS under HL at 29°C conditions (Scale bar = 1 mm). Quantification of real-time luminescence is below the kymographs where the red arrowhead denotes the downregulation of the *pIAA14::Luc* signal. The dashed grey rectangle in the kymographs mark the region for the real-time plots. The shaded area in real-time plots marks the signal originating from the root tip and local oscillation (root clock) zone.

**Supplementary Figure 20.**
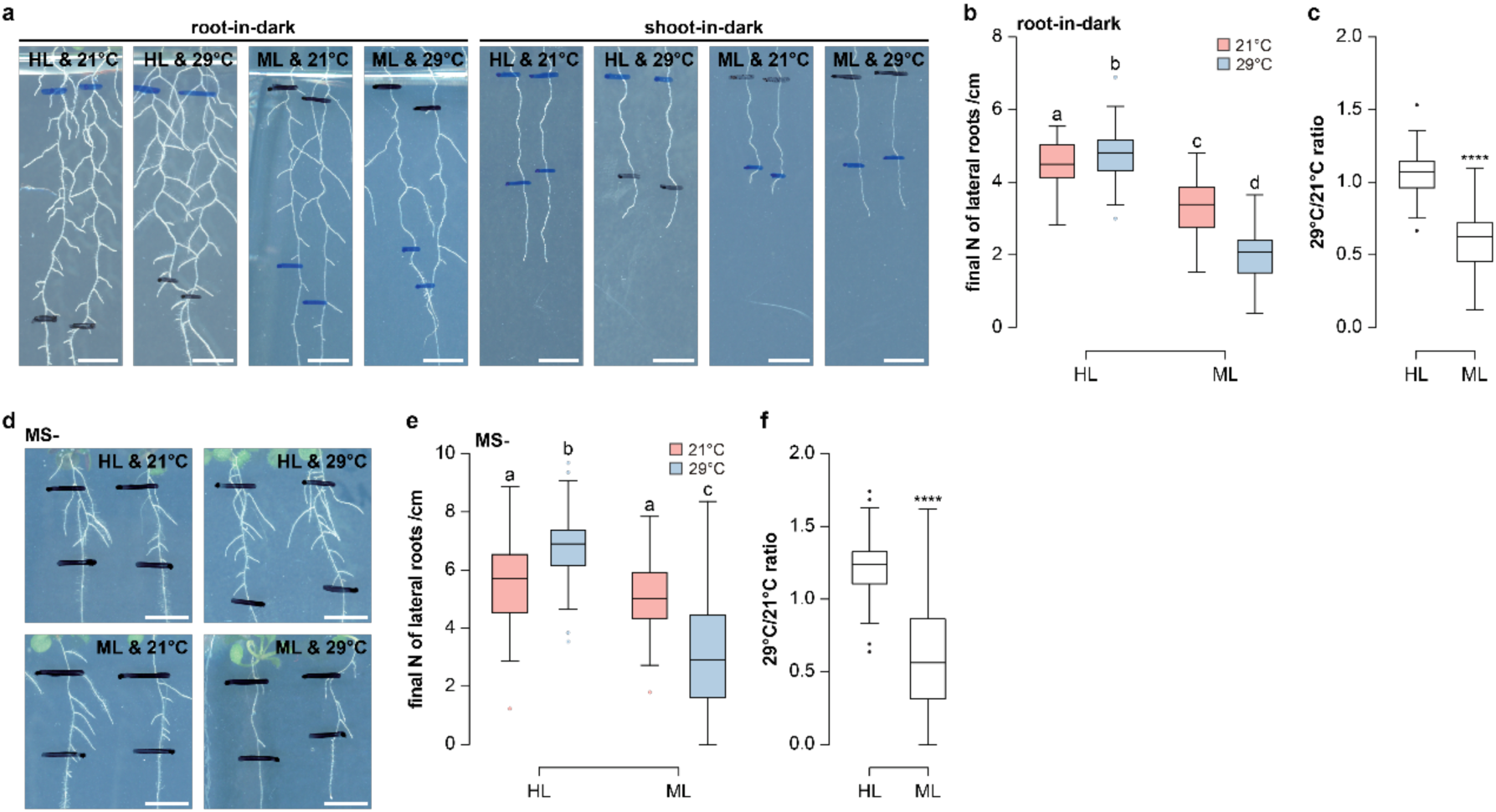
Shoot-received light modulates the high temperature-dependent inhibition of lateral root density while sucrose is mechanically distinct. **a**, Representative images of root systems at 21°C and 29°C under root-in-dark and shoot-in-dark conditions(Scale bar = 5 mm). **b**, **c**, Quantification (**b**) and calculated ratio (**c**) of LR density at 21°C and 29°C under root-in-dark conditions (*n* = 50 to 61). **d**, Representative images of root systems at 21°C and 29°C under high light (HL) and medium light (ML) conditions on solid medium without sucrose (MS-) plates (Scale bar = 5 mm). **e**, **f**, Quantification (**e**) and calculated ratio (**f**) of LR density of seedlings grown at 21°C and 29°C under HL and ML conditions on MS-plates (*n* = 47-51). Paired and two-tailed student’s t-test performed for (**c**) and (**f**) (*P <* 0.0001**** for both). Letters indicate values with statistically significant differences from one-way ANOVA performed for (**b**) and (**e**) (*P <* 0.0001 for both).

**Supplementary Figure 21.**
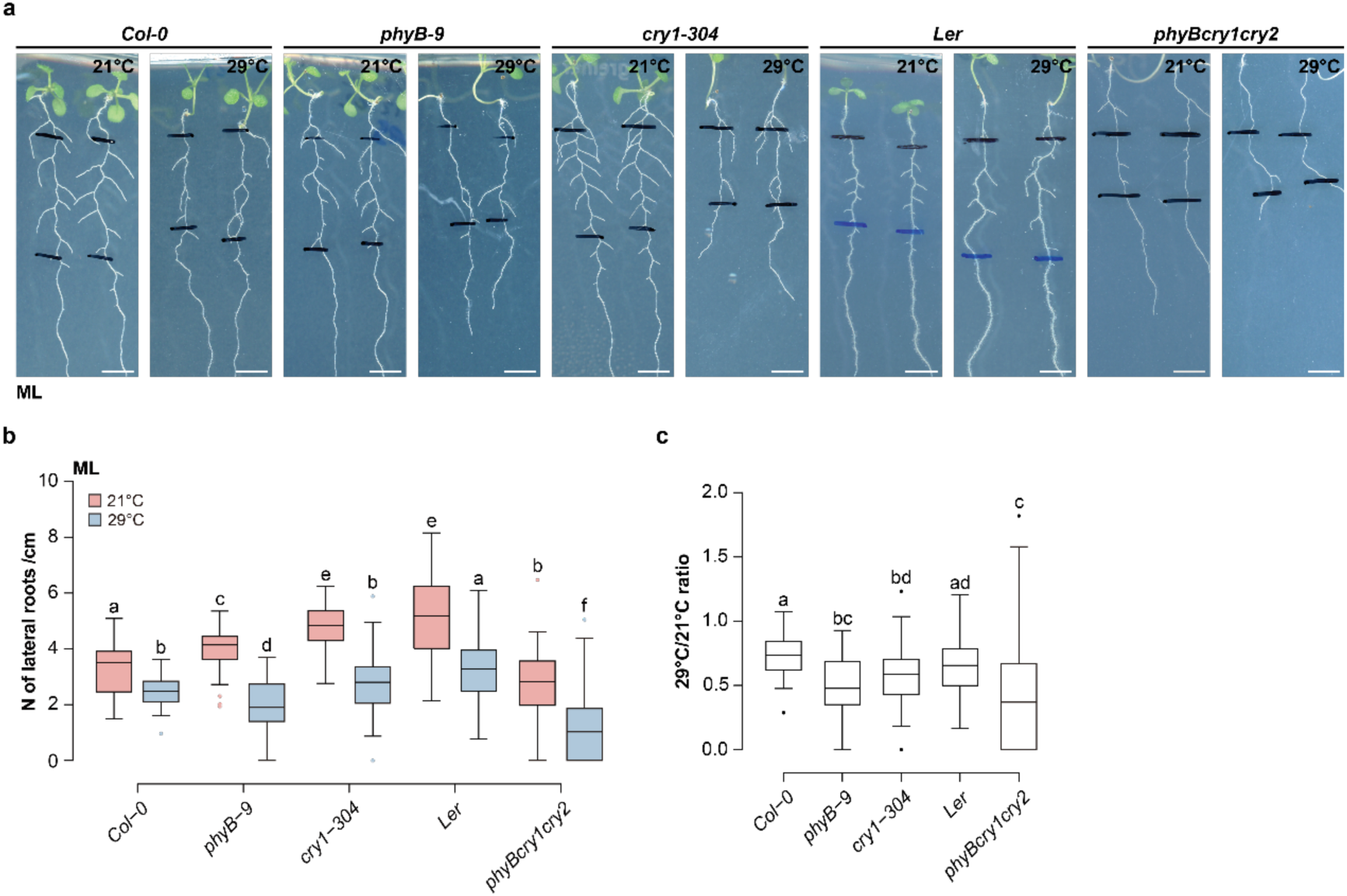
The loss of PHYB and CRY1 enhances the HT-dependent inhibition of LR under ML conditions. **a-c**, Representative root system images (**a**) and quantification (**b**) as well as the calculated ratio (**c**) of LR density in *Col-0*, *phyB-9*, *cry1-304*, *Ler* and *phyBcry1cry2* at 21°C and 29°C under medium light (ML) conditions (*n* = 51 to 90; Scale bar = 5 mm). Letters indicate values with statistically significant differences from one-way ANOVA performed for (**b**) and (**c**) (*P <* 0.0001 for both).

**Supplementary Figure 22.**
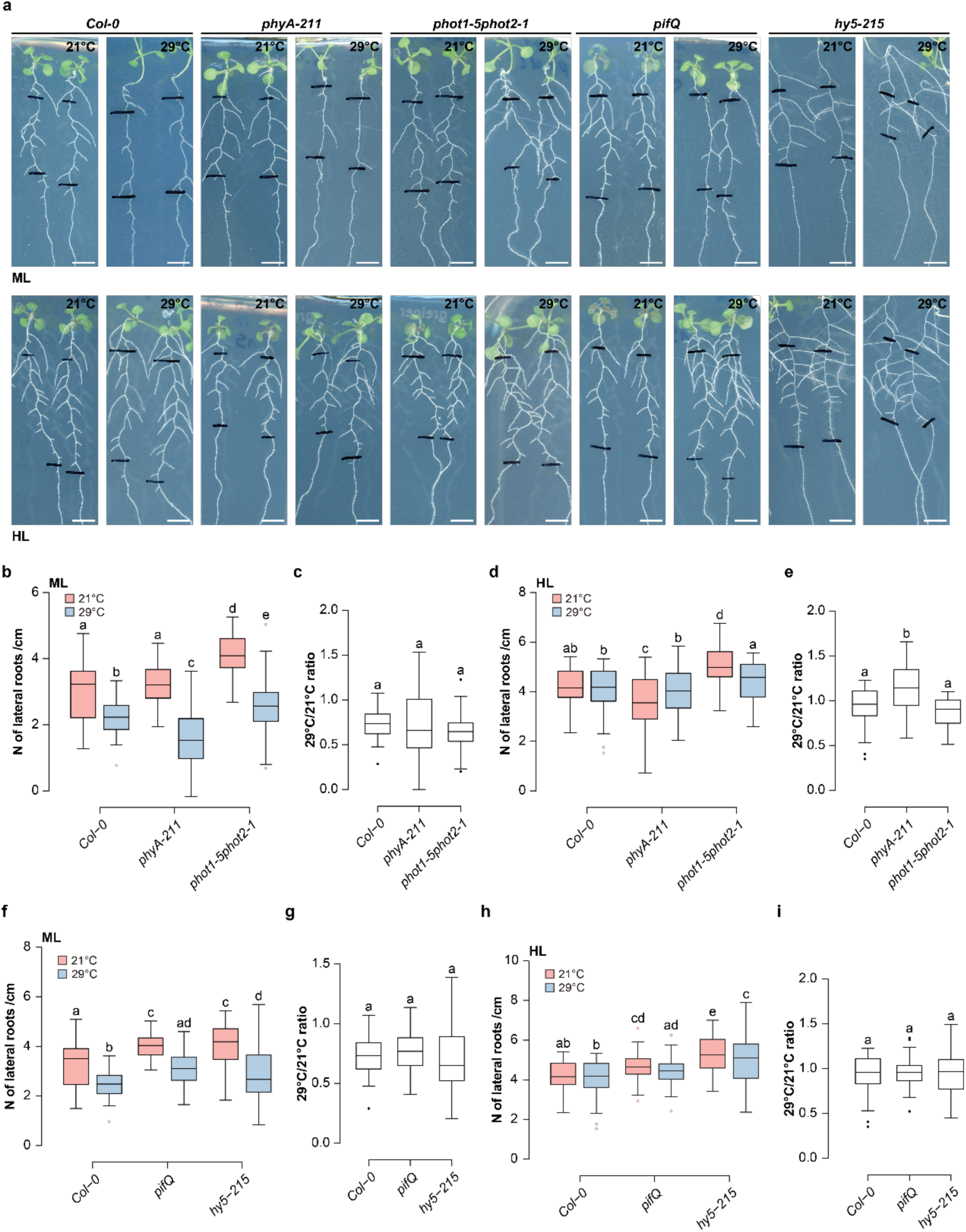
The inhibition of lateral rooting under high-temperature conditions in light signalling mutants. **a**, Representative root system images of wild-type *Col-0* and light signalling mutants at 21°C and 29°C under medium light (ML) (top row) and high light (HL) (bottom row) conditions (Scale bar = 5 mm). **b**, **c**, Quantification (**b**) and calculated ratio (**c**) of LR density in *Col-0*, *phyA-211* and *phot1-5phot2-1* at 21°C and 29°C under ML conditions (*n* = 45 to 54). **d**, **e**, Quantification (**d**) and calculated ratio (**e**) of LR density in *Col-0*, *phyA-211* and *phot1-5phot2-1* at 21°C and 29°C under HL conditions (*n* = 52 to 54). **f**, **g**, Quantification (**f**) and calculated ratio (**g**) of LR density in *Col-0*, *pifQ* and *hy5-215* at 21°C and 29°C under ML conditions (*n* = 52 to 54). **h**, **i**, Quantification (**h**) and calculated ratio (**i**) of LR density in *Col-0*, *pifQ* and *hy5-215* at 21°C and 29°C under HL conditions (*n* = 53 to 54). Letters indicate values with statistically significant differences from one-way ANOVA performed for (**b**), (**d**), (**e**), (**f**) and (**h**) (*P <* 0.0001 for all) and for (**c**), (**g**) and (**i**) (*P* > 0.05 for all).

